# The broccoli derivative sulforaphane extends lifespan by slowing the transcriptional aging clock

**DOI:** 10.1101/2025.05.11.653363

**Authors:** Christine A. Sedore, Erik Segerdell, Anna L. Coleman-Hulbert, Erik Johnson, Jonathan N. Levi, Gordon J. Lithgow, Monica Driscoll, Patrick C. Phillips

## Abstract

Sulforaphane, an organosulfur isothiocyanate derived from cruciferous vegetables, has been shown to inhibit inflammation, oxidative stress, and cancer cell growth. To explore the potential of sulforaphane as a candidate natural compound for promoting longevity more generally, we tested the dose and age-specific effects of sulforaphane on *C. elegans* longevity, finding that it can extend lifespan by more than 50% at the most efficacious doses, but that treatment must be initiated early in life to be effective. We then created a novel, gene-specific, transcriptional aging clock, which demonstrated that sulforaphane-treated individuals exhibited a “transcriptional age” that was approximately four days younger than age-matched controls, representing a nearly 20% reduction in biological age. The clearest transcriptional responses were detoxification pathways, which, together with the shape of the dose-response curve, indicates a likely hormetic response to sulforaphane. These results support the idea that robust longevity-extending interventions can act via global effects across the organism, as revealed by systems level changes in gene expression.

## Introduction

Without dramatic changes in the way that we approach the treatment of aging, it appears that recent gains in the increase in human lifespan are likely to slow over the next several decades [1]. At the same time, the exponential increase in the incidence of nearly every major disease late in life has led to a new and urgent focus on extending “healthspan” late in life rather than lifespan *per se*. This shift in attention in turn has led to an effort to identify drugs and other compounds that have positive impacts on health and longevity, rather than focusing on specific disease states [2]. Because of the long and costly effort required for new clinical drugs and the ongoing challenges around defining aging as a clinical disease state, previously approved drugs, such as rapamycin [3] and metformin [4], and natural product derivatives, such as resveratrol [5] and epigallocatechin-3-gallate (EGCG; found in green tea) [6–8] have led the way for clinical testing for potentially pro-longevity compounds. Three key elements for advancement of the geroprotection field are, first, the requirement that treatments lead to whole-body changes in aging trajectories at a functional level; second, a greater understanding of what the potential underlying functional targets may be; and, third, determination as to whether these compounds demonstrate effects at particular ages or whether they induce a steady accumulation of changes over the lifetime of an individual. Here, we use lifespan expansion and age-specific patterns of gene expression under treatment with the broccoli derivative sulforaphane to develop a unique “aging clock” within the nematode *Caenorhabditis elegans* to illustrate how these questions can be addressed using a widely available nutritional supplement.

Sulforaphane (SFN), an isothiocyanate found abundantly in cruciferous vegetables, was first isolated from broccoli by Zhang et al. [9], who found SFN to be a potent inducer of phase II detoxification enzymes. Numerous studies have since demonstrated that SFN possesses antioxidant, anti-inflammatory, and anti-cancer properties [10], with a particular focus on its induction of Nuclear Factor Erythroid 2-related factor 2 (NRF2), a major regulator of cellular detoxification responses and redox status [11,12]. Recently, SFN has also been shown to increase longevity in *C. elegans* by inhibiting DAF-2-mediated insulin/IGF signaling, promoting the nuclear localization of the FOXO transcription factor DAF-16, and thereby upregulating stress resistance and survival genes that include well-established regulators of stress resistance and longevity *sod-3*, *mtl-1*, and *gst-4* [13], [14]. SFN has also been shown to promote “healthy-aging” in *C. elegans* as measured by increases in mobility, pharyngeal pumping, and a decrease in lipofuscin accumulation [13], making it an ideal target for exploring translational potential more broadly.

A major development in aging research over the last decade has been the expansion of sets of molecular surrogates that change with age in a more or less clock-like manner. For example, so-called “epigenetic aging clocks” trained on DNA methylation profiles from whole-blood and tissue samples [15,16] can predict chronological age both within and between species fairly accurately. Newer aging clocks have been optimized to predict biological age using a variety of sophisticated, high-throughput approaches (e.g., transcriptomics, metabolomics [17–20]). Here we show that sulforaphane extends lifespan by more than 50% in a dose-dependent manner in *C. elegans* and that this intervention is required early in life to be effective. Whole genome transcriptomes gathered at multiple ages revealed a temporal shift toward “more youthful” expression of up to 20% in sulforaphane-treated individuals. Genes most strongly influenced by sulforaphane tend to exert their functions in compound detoxification and metabolism pathways, suggesting that sulforaphane may be enhancing longevity via a broad hormetic response instituted early in life that translates to enhanced health and longevity late in life. In addition to highlighting the specific benefits of sulforaphane, this work provides a general framework for the comprehensive application of robust and reproducible approaches, a hallmark of the *Caenorhabditis* Intervention Testing Program (CITP) [21], to aging clock research.

## Results

### Sulforaphane extends lifespan in a dose-dependent manner

We first sought to determine the dosage range for potential positive effects on lifespan in *C. elegans* using the standard laboratory strain N2 over six concentrations of sulforaphane (SFN) ranging from 10 to 800 µM (Fig. 1A and B). We found that SFN extended median lifespan at all but the highest concentration tested, with smaller effects on the low and high ends of the dosage range (12% increase in median lifespan at 10 and 400 µM, *p*<0.0001), peaking at 50 and 100 µM (53% increase, *p*<0.0001). We also observed a large increase in survival at 200 µM (41%, *p*<0.0001), with positive effects concentrated mostly later in life and therefore on average not as impactful as treatment with 100 µM SFN. Importantly, the positive effects of SFN abate at a concentration of 800 µM, especially in terms of mortality experienced later in life (*p*<0.0001) (Fig. 1A). Additionally, the 800 µM concentration of SFN resulted in a high percentage of censored individuals (50% vs 6% in controls), mostly due to vulval extrusion or individuals fleeing the plate. A bi-phasic dose-response curve is a classic hallmark of hormesis, in which small doses of a compound have positive effects on survival with higher doses becoming deleterious [22,23]. All subsequent experiments at this concentration of SFN, as it maximized overall lifespan extension.

**Figure 1:**
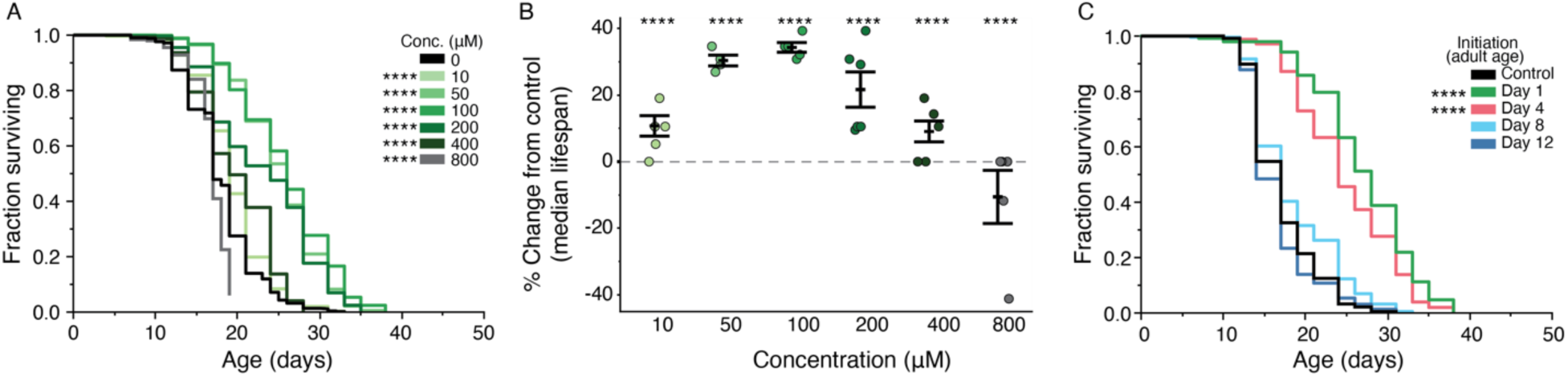
Sulforaphane extends lifespan in a dose-dependent manner with early life initiation of treatment. The effect of adult exposure to 10, 50, 100, 200, 400 and 800 µM sulforaphane on survival in *C. elegans* N2 beginning on day 1 of adulthood assessed via Kaplan-Meier curves of pooled replicates from at least two trials (A) and the percent difference in median lifespan from the control (B). For the latter, each point indicates the percent difference in the median lifespan of an individual trial plate relative to its specific control. Error bars represent the mean +/- the standard error of the mean. n = 859 (control) and 204-313 (SFN treated). C) Kaplan-Meier survival curves of survival after initiation of treatment with 100 µM sulforaphane at days 1, 4, 8, and 12 of adulthood. Day 1 initiation corresponds with our standard treatment protocol. Curves represent pooled replicates from two trials. n = 237-249. Asterisks represent *p*-values from the CPH model such that \*\*\*\**p*<.0001, \*\*\**p*<.001, \*\**p*<.01, and \**p*<.05.

### Sulforaphane treatment must be initiated early in life to enhance longevity

One question that often remains unanswered in terms of pro-longevity interventions is whether the treatment needs to be applied early in life to influence late-life survival, or whether intervention late in life is sufficient. Previous studies have shown interventions that work when administered early in life to have mixed outcomes when initiated at older ages [24–28]. We found that initiating treatment on both the first and fourth days of adulthood robustly and significantly extended lifespan (41-65% increase in median lifespan, *p*<0.001). However, beginning treatment later in life, on days eight and twelve of adulthood (close to when age-specific decline first begins to accelerate), failed to elicit a lifespan effect (Fig. 1C). Note that even these “late” ages occur well before animals reach the expected median lifespan. Consequently, we initiated sulforaphane treatment of the first day of adulthood for all subsequent experiments.

### Sulforaphane slows the transcriptional aging clock

To assess how SFN impacts the transcriptome during aging, we introduced SFN at adult day 1 and collected whole-animal RNA samples at five ages, ranging from day 2 of adulthood (24 hours post treatment initiation) until day 16 of adulthood (near median lifespan of the control) (Fig. 2A). Multi-dimensional scaling analysis across approximately 11,000 genes reveals an aging trajectory in which our older SFN-treated culture replicates more closely resemble their younger control counterparts late in life, with an average shift forward of approximately four days (Fig. 2A). That is, the day 2 and day 4 SFN replicates match closely with their day 2 and day 4 controls, respectively, but the day 12 SFN samples cluster with the day 8 controls, and the day 16 SFN cluster with the day 12 control replicates. This suggests an approximate 20% slowing of transcriptional aging at days 12 and 16 of adulthood.

**Figure 2:**
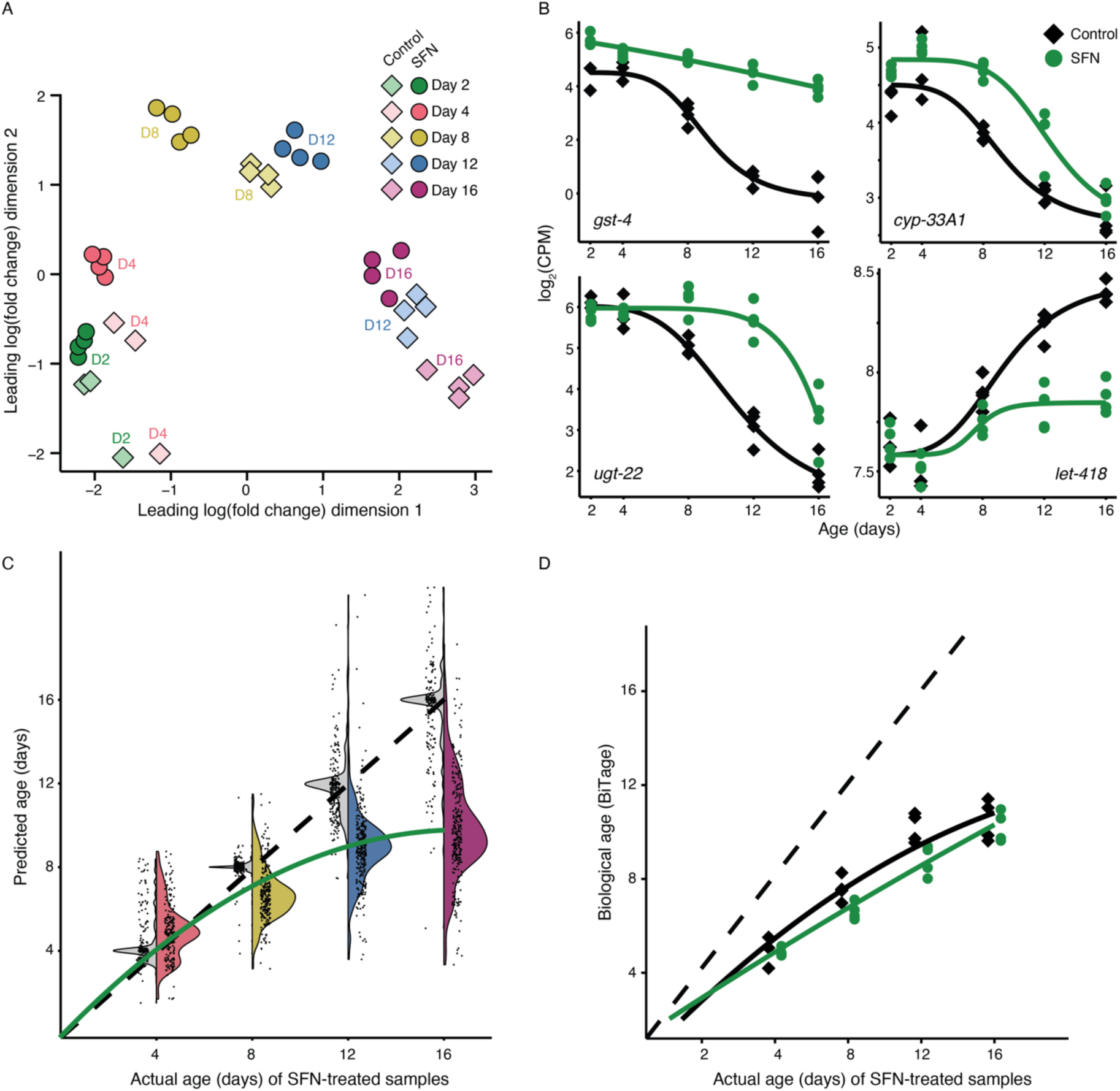
Sulforaphane slows transcriptional aging in *C. elegans*. A) Multi-dimensional scaling analysis of all RNA-seq replicates of *C. elegans* N2 treated with 100 uM SFN (circle) or DMSO control (diamond). Colors indicate the adult age at which the transcriptomic samples were collected: day 2 (green), day 4 (salmon), day 8 (gold), day 12 (blue), and day 16 (plum). Each point represents a single biological replicate. B) Gene expression trajectories *of gst-4, cyp-33A1*, *ugt-22*, and *let-418* in control (black lines) and SFN-treated (green) samples over age as exemplars for the full transcriptional clock gene set. Each point represents a single biological replicate. The normalized log_2_ counts per million (CPM) of each replicate are plotted over days 2, 4, 8, 12, and 16 of adulthood. Four-parameter logistic curves are shown for each control and SFN-treated series. C) Split violin plots showing the actual age of adulthood vs. the age predicted by our transcriptomic aging clock. Each point represents an individual gene from the clock and the black dashed line represents a 1-to-1 relationship between actual and predicted age. Gray violins correspond to the DMSO controls, and colors correspond to the adult ages at which the SFN treated samples were collected. The green line represents the best fit of the SFN samples. D) Biological ages predicted by BiT Age [18] are plotted over the actual age of our control and SFN-treated samples. BiT Age uses standardized chronological age, so the ages of the results are adjusted by two days to match our samples in adult age. Predicted ages of individual replicates (points are black for the controls, green for the treated samples) are plotted.

To more quantitatively assess the rate of slowing in transcriptional aging under SFN treatment, we sought to create an “aging clock” using the subset of genes displaying systematic changes in up or down regulation with age. To this end, we performed logistic regression of the expression trajectory of every gene in our control expression dataset and then used the resulting parametric model for the subset of genes displaying clear and reproducible age-specific changes (filtering for well-fitting models with *R*^2^ > 0.8) to create a gene-specific clock. The advantage of this approach over previous methods that produce a single composite age is that it allows an assessment of compound-specific effects on a gene-by-gene basis, as well as provides an appropriate null model for within-age variation within control samples (Fig. 2B & C). In total, our final clock consists of a subset of 311 genes (Extended Data Fig. 1). As null statistical model that captures the range of among-gene variation, the parametric age-specific clock generated a very close match between chronological and biological age for the control samples (Fig. 2C). For the SFN treated individuals, we were able to map the actual chronological age of SFN samples to the predicted age of the corresponding control with an equivalent level of expression on a gene-by-gene basis. Consistent with the whole genome analysis, the expression clock shows suppression of the expression trajectories for most genes in an age-specific fashion, with late-life expression patterns for SFN treated individuals matching expression of control individuals nearly half their age (Fig. 2C).

### Comparison to the BiT Age clock

Building on a composite of a large number of transcriptomic datasets for longevity extending mutants and other treatments in *C. elegans*, Meyer and Schumacher [18] constructed the BiT Age clock by normalizing these heterogenous datasets using elastic net regression to predict a single omnibus biological age from diverse RNA-seq samples. This analysis required temporal rescaling of each training sample and enforcing a standard median lifespan of µ = 15.5 days of adulthood. Further, the method reduces data complexity by eliminating quantitative details by binarizing the transcriptome in setting the value of a gene to 1 if its counts per million (CPM) is greater than the median CPM of the sample and a value of zero otherwise. These coefficients are summed across 576 predictor genes to yield a final prediction of biological age. For our samples, the BiT Age clock is successful in predicting that SFN treated individuals are somewhat “younger” than controls but fails in quantitative accuracy, with all samples—including controls—being predicted to be substantially younger than their actual chronological age (Fig. 2D). Interestingly, only one of the five genes specifically highlighted by Meyer and Schumacher [18] as being important drivers of the BitAge clock also appears in our clock: the intestinally produced and secreted innate immunity-related protein IRG-7 [29].

### The majority of differentially expressed genes are upregulated, with peak expression differences at day 12 of adulthood

To identify all potential genetic targets of SFN treatment, we also examined patterns of differential gene expression at each age separately across the entire dataset. Although treatment with SFN began on day 1 of adulthood, we see a substantial lag in the transcriptional response, with only 12 (0.13% of total) and 32 (0.34% of total) genes differentially expressed at days 2 and 4 of adulthood, respectively (Supplementary Table 1, Extended Data Fig. 2A and B). The overall response to SFN treatment reaches its maximum at day 12 of adulthood, with 1818 genes showing differential expression (14% of genes identified) (Fig. 3A). Of note, the balance of up and down regulation also changes with age, with between 80-100% of significantly impacted genes showing upregulation at ages 2, 4, 8, and 12, while there is a nearly equal balance of genes showing up and down regulation at day 16 (Extended Data Fig. 2D, Supplementary Table 1).

**Fig. 3:**
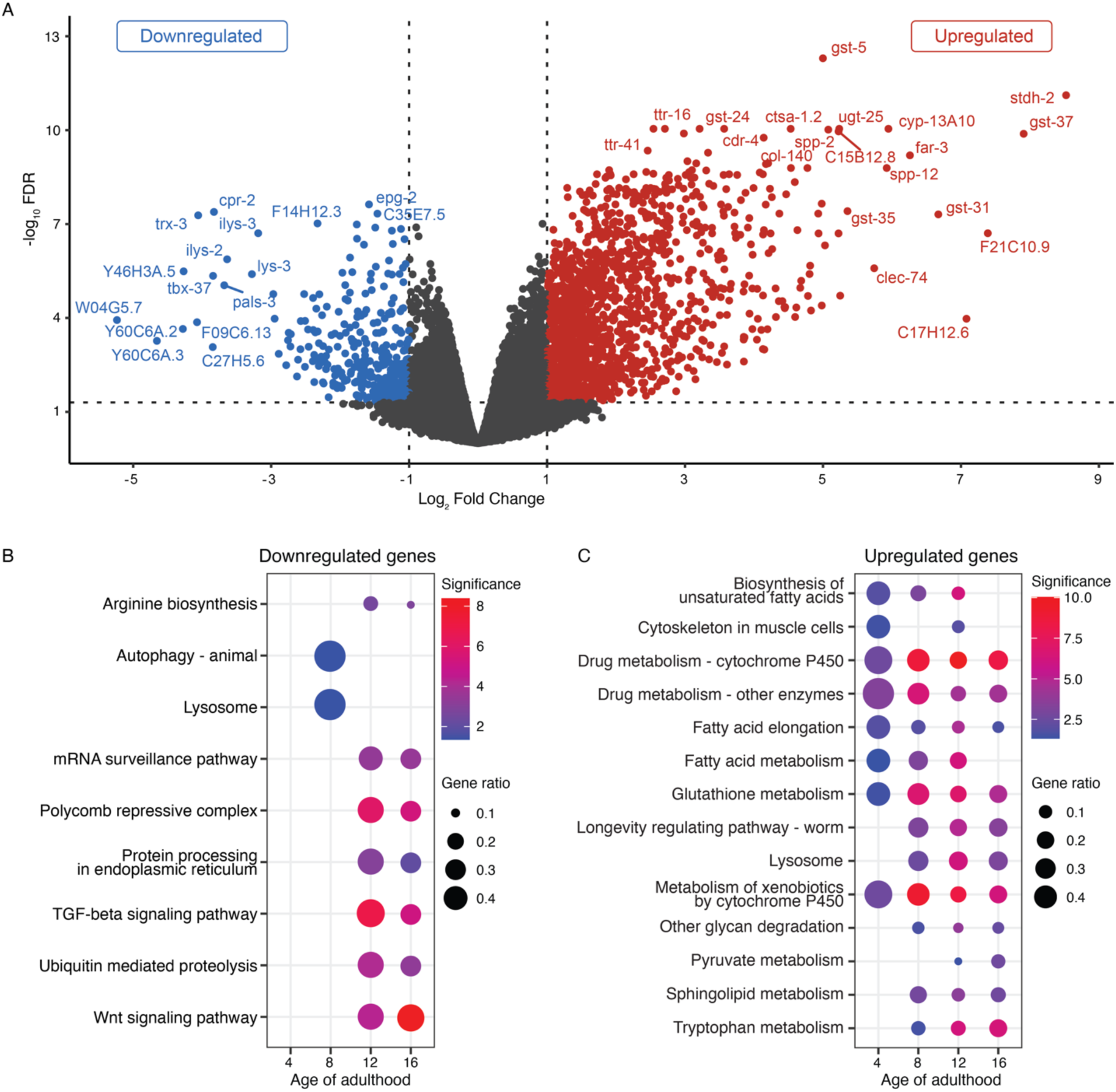
Transcriptional changes and KEGG enrichment for Longevity Pathways in *C. elegans* N2 at adult day 12. (**A**) Volcano plot of differentially expressed genes (DEGs) at day 12 of sulforaphane treatment. Significantly DE genes with absolute log_2_ fold change of at least 1 and FDR of 0.05 or lower are colored blue (downregulated) and red (upregulated). (B-C) KEGG (Kyoto Encyclopedia of Genes and Genomes; [32]) enrichment analysis for worm-specific longevity pathways by differential expression in total RNA. Dot plots show pathways which are enriched with B) downregulated and C) upregulated genes in animals aged 4, 8, 12, and 16 days of adulthood. Point size corresponds to the fraction of genes (gene ratio) in the KEGG category that are differentially expressed. The significance of a category at each age is calculated by taking the negative log10 of the adjusted p-value; the color scale changes from blue to red, with red indicating greater significance. Volcano plots of all other ages are in Extended data figure 2.

### Targeting of detoxification and metabolism pathways

Among the upregulated genes, we find significant enrichment of genes associated with detoxification. At day 12, several glutathione-s-transferases are among the genes with the highest fold-change, along with cytochrome P450s and uridine diphosphate-glycosyltransferases (Fig. 3A), with this general pattern remaining true across all ages. Both UDP-glycosyltransferases and glutathione-s-transferases are phase II detoxification enzymes [30]. Sulforaphane and its analogues have previously been identified as potent inducers of phase II detoxification enzymes, including glutathione transfer activities [9,31]. To examine pathway-specific responses more comprehensively, we decided to explore which pathways may be involved in this response with Kyoto Encyclopedia of Genes and Genomes (KEGG) [32], Gene Ontology (GO) [33,34], and Reactome [35,36] enrichments (Fig. 3B and C, Extended Data Fig. 3). As expected, the KEGG worm-specific longevity promoting pathways upregulated at all ages were detoxification pathways, including drug and xenobiotic metabolism and glutathione metabolism. Multiple fatty acid pathways were upregulated: biosynthesis of unsaturated fatty acids, along with fatty acid elongation and metabolism (Fig. 3B and C). Other metabolism pathways upregulated at older ages include pyruvate, tryptophan, and sphingolipid metabolism.

We also performed KEGG enrichment analysis on a subset of the 311 aging clock genes and found that fatty acid metabolism, fatty acid degradation, and glutathione metabolism are the main pathways associated with general drivers of aging (*p*=0.037 for each category). Looking at just the 10% or 20% of genes with the lowest predicted ages in the clock at days 12 and 16 revealed no significant KEGG enrichment categories, indicating that there is no clear thematic structure to the function of the clock genes.

Because our KEGG analysis reinforced the view that SFN was resulting in a detoxification response, we examined global effects across key detoxification gene families more specifically. In drug metabolism, cytochrome P450 (CYP) enzymes are involved in phase I detoxification, while UDP-glycosyltransferases (UGTs) and glutathione-s-transferases (GSTs) carry out phase II detoxification [37]. In general, the CYPs and GSTs were significantly upregulated at every age (Fig. 4; Supplementary Table 2). Similarly, the UGTs were significantly enriched as well starting at day 8. This difference is particularly striking at day 16, where 6% of total genes were upregulated (defined as >1 abs(logFC) and <.05 FDR), while 57-72% of all CYPs, GSTs, and UGTs identified were upregulated (*p*<1.0×10^-13^).

**Figure 4:**
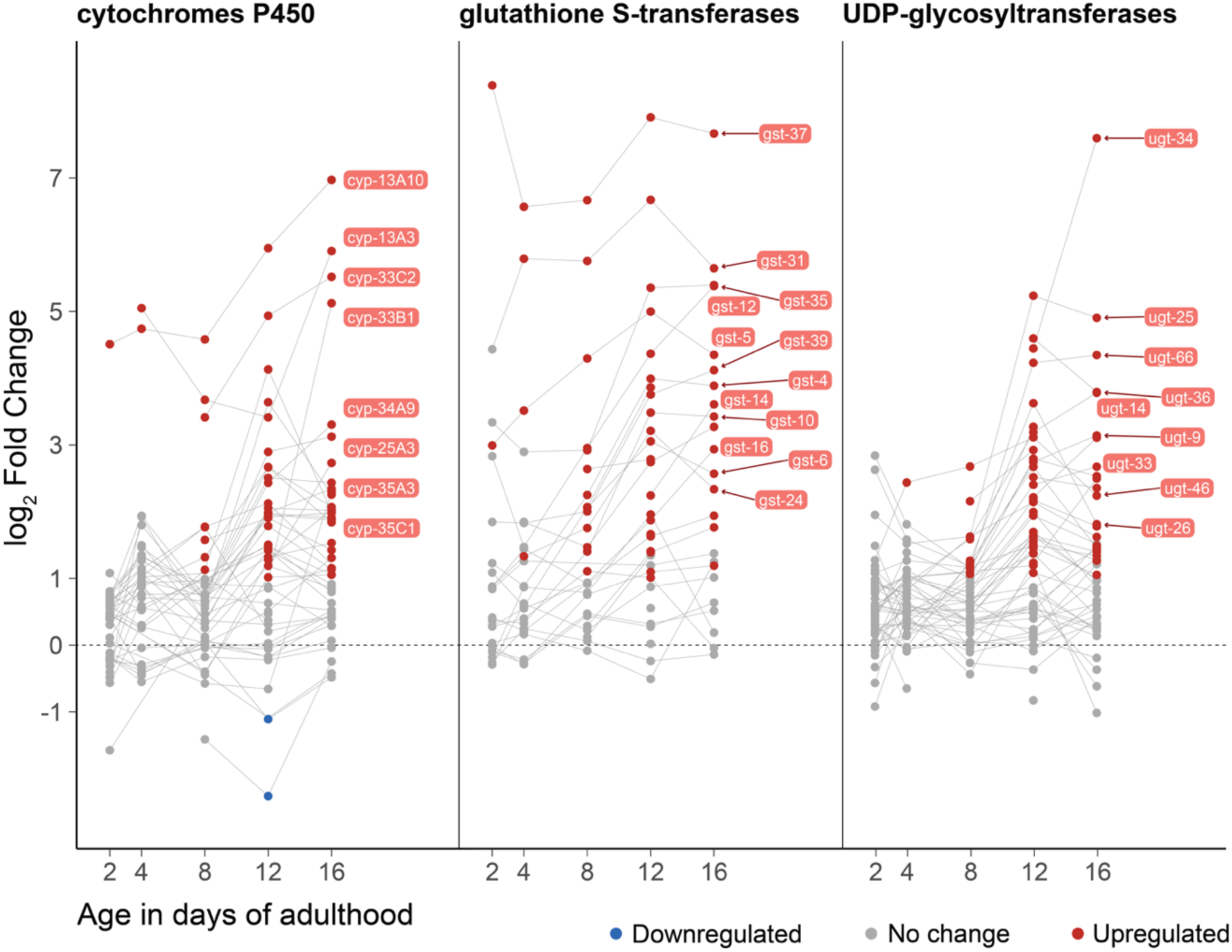
Age-specific changes in expression for detoxification-related gene families. Effect of SFN treatment on CYP, GST and UDP genes at days 2, 4, 8, 12, and 16 of adulthood. Dots represent gene expression levels at each age; colors indicate upregulation (red), downregulation (blue), or no change (gray). Lines connect each gene between ages; genes with a log_2_ fold change greater than 3 at any age are labeled. Expression data for these genes is available in Supplementary information table 3; analysis of enrichment for these genes is in Supplementary information table 6; a list of all genes in this figure is available as Supplementary information table 7.

### Overlap of genes differentially expressed after sulforaphane treatment and published *skn-1* and *daf-16* expression datasets

Because previous observations highlighted dependency the SFN response on the DAF-16/FOXO transcription factor for lifespan extension in *C. elegans* [13] and because SFN is a known activator of the Nrf2/SKN-1 transcription factor [38,39], we explored the overlap of genes known to be targeted by DAF-16 and SKN-1 and the genes displaying differential expression in our samples (Supplemental Data Fig. 4), as measured by RNA-sequencing [40–42] or microarrays [14,43,44]. We find variable overlap of SKN-1 expression datasets with our SFN expression results, with Oliveira et al. [43] dataset ranging from 18-35% overlap, but the other two studies falling in the 4-16% range. It should be noted that most of these studies examined changes in expression of SKN-1 targets under in the absence of experimental stressors, which may explain the lack of overlap. However, Oliveira et al. [43] also looked at SKN-1 upregulated genes under both arsenite (As) and t-BOOH stress, which we compared to both SFN upregulated and SKN-1 upregulated genes under no-stress conditions. Notably, the overlap of SFN results with the SKN-1 the datasets did not differ whether samples where under stress or not—21-38% overlap under no stress, 26-37% overlap under stress (with the exception of day 8 SFN and SKN-1 under t-BOOH stress, which showed only 6% overlap). This is perhaps unsurprising as the majority of As-upregulated genes are also upregulated under no-stress conditions, and only a small subset of genes upregulated under t-BOOH stress are SKN-1 dependent.

When comparing published DAF-16 expression profiles with our SFN datasets, we again see some small overlap, particularly at ages 12 and 16. Three of the four tested datasets show >20% overlap with our SFN samples at those ages, while the remaining dataset only overlaps 12-17%. In general, there is minimal overlap among any of the compared datasets, with none of the independent studies sharing a majority of genes. This highlights challenges of reproducing expression studies conducted under differing experimental conditions and underscores the value of the CITP, in which all experiments are performed under conditions of rigorous standardization [21].

## Discussion

Aging is a complex process characterized by the progressive deterioration of physiological function over time. Efforts to understand and ameliorate age-related decline have been ongoing for several decades, with numerous studies demonstrating the ability to extend lifespan and healthspan in a variety of organisms through specific interventions including dietary restriction, exercise, and genetic manipulation [45–48]. The identification of lifespan-enhancing small molecule compounds has likewise emerged as an important intervention, with recent studies demonstrating the ability of single compounds to increase longevity and extend healthspan [49–52]. To this end, the CITP was created to utilize the model organism *C. elegans* for this purpose, a model that has proven invaluable in the initial screening and characterization of such compounds due to the relatively short lifespan, low cost, and ease of experimental manipulation [53–55]. Based on previous findings and the recommendation from the CITP Access Panel, in which the research community as a whole can suggest compounds for investigation, we tested the broccoli derivative sulforaphane for lifespan intervention in wildtype *C. elegans* as a “sponsored compound” through the CITP pipeline [21].

Sulforaphane is an organosulfur isothiocyanate derived from the glucosinolate precursor, glucoraphanin. Glucosinolates are a family of plant-derived compounds found abundantly in broccoli (*Brassica oleracea*) and other cruciferous vegetables. Upon damage to the plant (e.g., from cutting, chewing, or infection), the plant-defense enzyme myrosinase increases and hydrolyzes glucoraphanin to produce sulforaphane [56]. While many epidemiological studies have linked fruit and vegetable intake to improvements in health outcomes, it may be worth noting that cruciferous vegetable intake in middle age has been associated with lower risk of all-cause mortality [57]. Several clinical studies have demonstrated positive health effects from sulforaphane treatment, including: the reduction of fasting blood glucose and HbA1c in patients with type-2 diabetes [58]; the reduction of carcinogen-induced oral cancer in mice [59]; the prevention of cartilage destruction in humans (both *in vitro* and *in vivo*), with potential for the treatment of osteoarthritis [60]; and increases in levels of the antioxidant glutathione in healthy human subjects, with implications for the treatment and management of neurodegenerative and neuropsychiatric disorders [61]. As cancer, diabetes, osteoarthritis, inflammation, and neurodegenerative diseases are all associated with aging and age-dependent decline, these trials, when taken together, indicate a potentially significant role for sulforaphane in generalized targeting of the pathways that promote healthy aging and delay mortality.

This view is also supported by the fact that sulforaphane targets multiple hallmarks of aging [62] by reducing inflammation, impacting epigenetic regulators, activating proteasome components, promoting stem cell proliferation *in vitro*, and preventing cell senescence *in vitro* [63–66]. In mice, the prevention by sulforaphane of diabetic cardiomyopathy was associated with upregulation of NFE2-related factor 2 (Nrf2) expression and an increase in the expression of Nrf2-dependent downstream antioxidants [38]. Additional studies have shown that sulforaphane also increases Nrf2 expression in other mice tissues [39]. Nrf2/SKN-1 is involved in detoxication and antioxidant defense, among other functions. Taken together, these data suggest important age-related pathways impacted by sulforaphane treatment and highlight interest in further evaluation of the role of sulforaphane in mediating lifespan and healthspan.

We demonstrate that sulforaphane extends lifespan in *C. elegans* by more than 50% and that such lifespan intervention must be administered early in adult life (days 1 or 4 of adulthood) rather than late/pre-senescence (days 8 and 12 of adulthood), which is consistent with earlier work [13]. Despite early adult intervention being required for lifespan extension via sulforaphane, whole-organism gene expression does not change until later in life and peaks on day 12 of adulthood (midlife for treated animals), with roughly 14% of the genome responding to sulforaphane treatment. Moreover, the majority of differentially expressed genes are upregulated by 53-100% at all ages measured. This suggests a regulatory cascade with a long temporal trigger and/or a need to accumulate sufficient SFN to activate a response. Interestingly, though sulforaphane has been shown to activate Nrf2/SKN-1 and requires FOXO/DAF-16 activity, we found only marginal overlap between our dataset (up to 35% for the former and 37% for the latter) and existing expression profiles for mutants of each, but also that these studies also did not replicate one another. This result is likely driven by differences in methodology across studies which, in turn, highlights the value and necessity of highly consistent and replicated studies of transcription in aging studies. In contrast, we find a very clear and consistent signal of large-scale upregulation of genes involved in detoxification pathways, particularly at later ages. This reinforces the observation of dose-dependent effects of sulforaphane on longevity and suggests that sulforaphane slows the aging clock via hormetic induction of stress responses at low doses.

There is increasing evidence supporting the enrichment in the activity of detoxification genes in long-lived individuals across a variety of organisms [67]. A meta-analysis looking at the common gene signatures of longevity interventions found that the transcription levels of detoxification enzymes cytochrome P450s (CYPs) and glutathione-*S*-transferases (GSTs) were enriched in mice livers across interventions [68]. For instance, Tyshkovskiy et al. [68] showed that several glutathione S-transferase (GST) genes were significantly upregulated across multiple interventions (which included genetic, pharmacological, and dietary interventions), including *Gstt2*, *Gsto1*, and *Gsta4*, and further found that the most significantly upregulated functions across interventions included drug metabolism by cytochrome P450 and glutathione metabolism activated by the NRF2 pathway. Likewise, a multi-level cross-species comparative analysis in mutant nematodes, fruit flies and mice with reduced insulin/IGF-1 found that lifespan extension was correlated with significantly higher transcript levels of genes encoding certain detoxification enzymes, including GSTs and cytochromes P450 (increased in long-lived worms and flies, but only slightly in mice) [69]. These detoxification gene classes have long been implicated for their role in aging in *C. elegans* [70]. We previously observed that lifespan thioflavin T extends lifespan across Caenorhabditis strains and species and also requires NRF2/SKN-1 function in N2 [50,71]. Recent studies in mice [72] and nematodes have also shown that pharmacological interventions capable of extending longevity are associated with an increase in these drug metabolism/detoxification pathways [73–75], suggesting that targeting detoxification *per se* may be a useful longevity intervention therapy, concordant with our results presented here.

### A novel aging clock

We used a subset of genes with strong age-dependent patterns of expression to create a clock based on “normal” aging, finding that treatment with sulforaphane results in a ∼20% delay in apparent biological aging as measured by transcriptional abundance. This nicely echoes the benefits of sulforaphane treatment on overall longevity. Previous transcriptome-based aging clocks such as BiT Age have been designed to dissociate chronological from biological age using heterogeneous datasets to create a single omnibus estimate of biological age [18]. By contrast, we aimed to develop a clock that would enable us to assess and visualize aging effects on a gene-by-gene basis. Perhaps not surprisingly, our tuned aged clock outperforms BiT Age for our samples but also provides an estimate for the among gene variation in reproducibility within our control samples as well, which allows a more robust statistical framework for assessing clock quality and reproducibility. Further, gene-specific changes allow us to understand how the clock genes themselves are affected by sulforaphane. In this case, pathway-specific changes in the clock genes are very similar to the patterns revealed by age-specific analyses across the entire transcriptome. Age trajectory approaches are likely to reveal much more detail in transcriptional aging than simple two-point analyses in general [76]. It is nevertheless important to remember that all aging clocks basically reveal biomarkers of aging and cannot directly be interpreted as revealing underlying causes of aging [77,78].

Further, these analyses are based on whole-animal transcriptomics, whereas other studies have found significant tissue-specific heterogeneity across other measures, such as the epigenetic clock [79]. There are also several potential statistical issues that deserve greater attention in the future, although they do not appear to affect our particular results. First, it is well known that aging clocks can be susceptible to sampling variation [77]. and so more in-depth investigation of potential sources of variance in the baseline, control clock is merited. Second, particularly high levels of gene expression in treated samples have the potential to “overrun” the clock, making it impossible to calculate an age-matched estimate through the model and thereby causing a potential downward bias in apparent age in treated samples, as only expression values that fall within the range of control values can be retained in the clock estimates. This happens very rarely for our sulforaphane samples and is therefore not a source of bias in the values presented here, but this could be an issue for application to other compounds and bears further investigation, especially since total variation in expression late in life is likely both to be a hallmark of aging and to be a statistical challenge for accurate estimation [80]. Overall, while there clearly remains a great deal to be explored in this arena, our approach reveals that significant insights into the functional basis of the pro-longevity effects of compounds can be made through the rigorous development of whole-transcriptome clocks.

### Conclusion

Although the exact translatability of these results to human populations remains to be seen, the general admonition that appears to emerge from this work is: Eat your broccoli, just not too much! The large body of mounting evidence of the positive effects of sulforaphane on many health-related traits in humans suggests that direct translation to longevity *per se* is quite likely. Sulforaphane is usually described as a Nrf2 agonist [81], and may be directly acting as a direct modifier of this pathway, which would recommend it as a particularly useful pro-longevity intervention. However, *C. elegans* lacks a homolog to the Keap1 Kelch-like ECH-associated protein, which is thought to be the actual target of sulforaphane [82], indicating that an alternate mode of action must be involved in this case. Alternatively, the hormetic, stress-pathway inducing properties of sulforaphane may indicate that many beneficial dietary supplements work in a fairly generic fashion as mild toxins rather than being driven by the biochemical properties of the compounds themselves (e.g., as antioxidants). This idea is best addressed by comprehensive pathway-specific analysis of responses across a wide-ranging set of pro-longevity compounds, such as has been done for sulforaphane here.

## Methods

### Nematode strain and maintenance

*C. elegans* N2_PD1073 was obtained from the *Caenorhabditis* Genetics Center. This strain is a clonal line derived from the N2 strain VC2010 that was used in the generation of a new N2 reference (“VC2010-1.0”) genome [83]. Strains were maintained at 20°C and 80% RH on 60 mm NGM plates seeded with *E. coli* OP50-1.

### Compound intervention

Compound intervention was conducted as previously published [71,84,85]. Sulforaphane (D,L-Sulforaphane, Santa Cruz Biotech, sc-207495D) was mixed with DMSO (dimethyl sulfoxide) to create a stock solution. Stock solutions (DMSO for the controls) were then diluted with water to create working solutions and distributed onto 35 mm NGM plates containing 51 µM FUdR and previously seeded with OP50-1. Plates were treated with compound solutions such that the final concentration was calculated given the volume of NGM per plate and the final concentration of DMSO was 0.25%.

### Lifespan assays

Manual lifespan assays were performed as described [71,84,85]. Briefly, populations were reproductively synchronized via timed egg lays on 60 mm NGM plates. On day one of adulthood, animals were transferred (50 individuals per plate) to 35 mm NGM plates containing 51 µM FUdR and compound intervention (sulforaphane or solvent control). Individuals were transferred again to fresh plates on days two and five of adulthood, and once weekly thereafter. Each individual was examined three times per week and defined as dead if no response was observed after stimulation with a platinum pick. Animals were censored for internal hatching of embryos, vulval extrusions, or if they escaped the plate.

### Late life intervention

Late life intervention was conducted as described above, with these changes: All individuals were transferred to solvent control plates on day one of adulthood following the regular procedure for control populations above, with the exception of day one initiation cohort, which was transferred to 100 µM sulforaphane plates as indicated for the treatment populations above. Worms were then transferred on days two, four, eight, and twelve of adulthood from the control plates to correspond with initiation of sulforaphane treatment of one of the cohorts. Each cohort was maintained and scored on control plates until initiation of treatment. After treatment was initiated on all cohorts, animals were transferred weekly to fresh plates until death.

### Statistical analysis of survival

Statistical analyses of lifespan experiments were conducted following standardized CITP analysis pipeline [71]. Briefly, a mixed-model approach was used where compound/concentration was treated as a fixed effect, and other potential variables (start date, experimenter, and plate replicate) were considered random effects. Survival was analyzed in R ([86]; v4.2.3 or 4.3.3) both with a generalized linear model using the lme4 package (v1.1-32 or v1.1-35), and with a mixed-effects Cox-Proportional Hazards (CPH) model with the coxme package ([87]; v2.2-18.1 or v2.2-20). Analysis was done within a strain to allow for each compound intervention to be compared to its specific control in the randomized blocks design. Each compound effect was determined via a planned comparison between each compound/concentration and its control. Statistics reported in figures are from the CPH model.

### Transcriptomics

For RNA-sequencing, animals were synchronized and compound treated as described above. Four biological replicates were aged and collected in tandem, with approximately 50 worms total per age, intervention, and replicate. Worms were picked into 0.2 mL tubes containing 45 µL DNA elution buffer (Zymo) and 5 µL proteinase K (50 µL total), then flash frozen with liquid nitrogen. Libraries were prepared using the KAPA mRNA HyperPrep kit (Kapa Biosystems, KK8580) as per manufacturer protocol except that the total volume was adjusted to ¼ per reaction. Final libraries were normalized by concentration and sequenced on an Illumina Novaseq 6000 with the SP 100 cycle (Genomics and Cell Characterization Core Facility, University of Oregon).

Single-end FASTQ files for all sulforaphane treated and DMSO control samples were aligned to the *C. elegans* WBcel235 (build 108) reference genome using the Subread package [88] version 2.0.2. Uniquely mapped reads were assigned to *C. elegans* genes with the featureCounts program [89]. Subsequent filtering, normalization, and differential expression analysis were performed with the edgeR package [90] version 3.28.1, using R version 3.6.2. Lowly expressed genes were removed from each dataset; only genes that had at least 10 reads in at least four samples and a minimum total count of 15 reads across samples were retained. To remove composition biases between libraries, the library sizes were normalized using a trimmed mean of M-values (TMM) between each pair of samples [91]. A pairwise expression analysis was performed on the transcriptomes of the treatment and control samples from worms aged 2, 4, 8, 12, and 16 days of adulthood. Quasi-likelihood (QL) F-tests for treatment vs. control sample effect were carried out on fitted gene-wise negative binomial generalized log-linear models. P-values were corrected for false discovery using the Benjamini-Hochberg method. Genes were considered significantly differentially expressed at an absolute log2 fold change of at least 1 and an FDR of 0.05 or lower.

Venn diagrams comparing the differentially expressed genes from our dataset to previously published expression datasets were made in R using the VennDiagram package (v1.7.3). Gene names (or sequence names if provided) were converted into their Wormbase ID using the Gene Name Sanitizer tool on Wormbase (https://wormbase.org/tools/mine/gene_sanitizer.cgi, accessed 5/2025) if not already provided. Obsolete genes or genes not found in Wormbase were excluded from analysis. Duplicates were removed from all datasets before Venn diagrams were generated.

### Pathway and ontology enrichments

Gene ontology and pathway enrichment analysis was performed in R with the clusterProfiler ([92] v.14.3) and ReactomePA ([36] v1.30.0) packages. Gene Ontology biological process and molecular function categories [33,34], KEGG (Kyoto Encyclopedia of Genes and Genomes) [32] and Reactome [35] pathways were tested for over-representation in the set of differentially expressed genes at days 2, 4, 8, 12, and 16 of adulthood. *P*-values were corrected for false discovery using the Benjamini-Hochberg method and categories were considered significant at an FDR of 0.05 or lower.

### Transcriptional aging clock

To build the transcriptional aging clock, we performed a systematic logistic analysis of the expression trajectory of every gene in the RNA-sequencing dataset, fitting parametric models to the normalized log-CPM values of control samples over ages 2, 4, 8, 12, and 16 days of adulthood, and selecting genes with non-zero slope that exhibited a statistically significant temporal change in expression. The four parameters of the logistic model are the (*a*) minimum and (*d*) maximum values that can be obtained, the (*c*) point of inflection (the point on the S-shaped curve midway between the minimum and maximum), and the (*b*) Hill’s slope of the curve (related to the steepness of the curve at the inflection point). Parameters for each gene were calculated with the R package dr4pl (v2.0.0). A coefficient of determination (R-squared) was calculated for each of the selected gene-specific models; those with an R-squared value greater than 0.8 were selected for the final clock, a total of 311 genes. Expression levels of the same 311 genes in sulforaphane-treated samples at each biological age were used to predict equivalent ages in non-treated worms on a gene-by-gene basis. Using the parameters estimated for each gene (*a*, *b*, *c*, and *d*) and the log-CPM expression in SFN-treated samples (*y*), the predicted age (*x*) is calculated with the following formula:

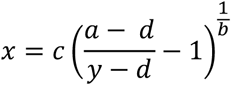

Biological ages of control and treated RNA-sequencing samples at ages 4, 8, 12, and 16 days of adulthood were also predicted using the BiT Age clock ([18]; https://github.com/Meyer-DH/AgingClock). The prediction function took as input (A) our count-per-million normalized RNA-seq counts with genes as rows and samples as columns and (B) the predictor genes and corresponding regression coefficients in the BiT Age model. The WormBase IDs of three genes (WBGene00016086, WBGene00015370, and WBGene00007532) were revised in the BiT Age table following the merging of gene records in WormBase, making IDs fully consistent with those used in our dataset. The predicted ages were then used to calculate a second biological age correction as described by the BiT Age authors. We ran a separate analysis using the mean CPM gene expression values across replicates at age in each treatment group.

## Acknowledgments

This work was supported by funding from National Institutes of Health grants (U01 AG045844, U01 AG045864, U01 AG045829, U24 AG056052). The nematode strain used here was provided by the *Caenorhabditis* Genetics Center, which is funded by NIH Office of Research Infrastructure Programs (P40 OD010440). We thank Tiziana Cogliati, Max Guo, and Viviana Perez of the NIA, Daniel Promislow, and members of the Phillips, Driscoll, and Lithgow labs for helpful discussions. Additionally, we acknowledge members of the CITP Steering Committee and Access Panel for their feedback and engagement.

## Data availability

Full CITP standard operating procedures can be accessed at CITPaging.org.

Lifespan data is available at CITPaging.org/portal and in Source Data Fig. 1A and 1B and Source Data Fig. 1C. Relevant R scripts can be found in Supplementary Software R_scripts.zip. Transcriptomic data have been deposited in NCBI’s Gene Expression Omnibus (Edgar et al., 2002) and are accessible through GEO Series accession number GSE289233 (https://www.ncbi.nlm.nih.gov/geo/query/acc.cgi?acc=GSE289233).

---

**Extended Data Fig. 1:** Expression trajectories of the complete set of 311 aging clock genes in control (black lines) and SFN-treated (green) samples over age. Each dot represents a single biological replicate. The normalized log_2_ counts per million (CPM) of each replicate are plotted over days 2, 4, 8, 12, and 16 of adulthood. Four-parameter logistic curves are shown for each control and SFN-treated series; logistic curves are omitted from SFN-treated series where parameters could not be computed for the given gene. ExtendedData1_Trajectories_of_all_clock_genes.pdf

**Extended Data Fig. 2:**
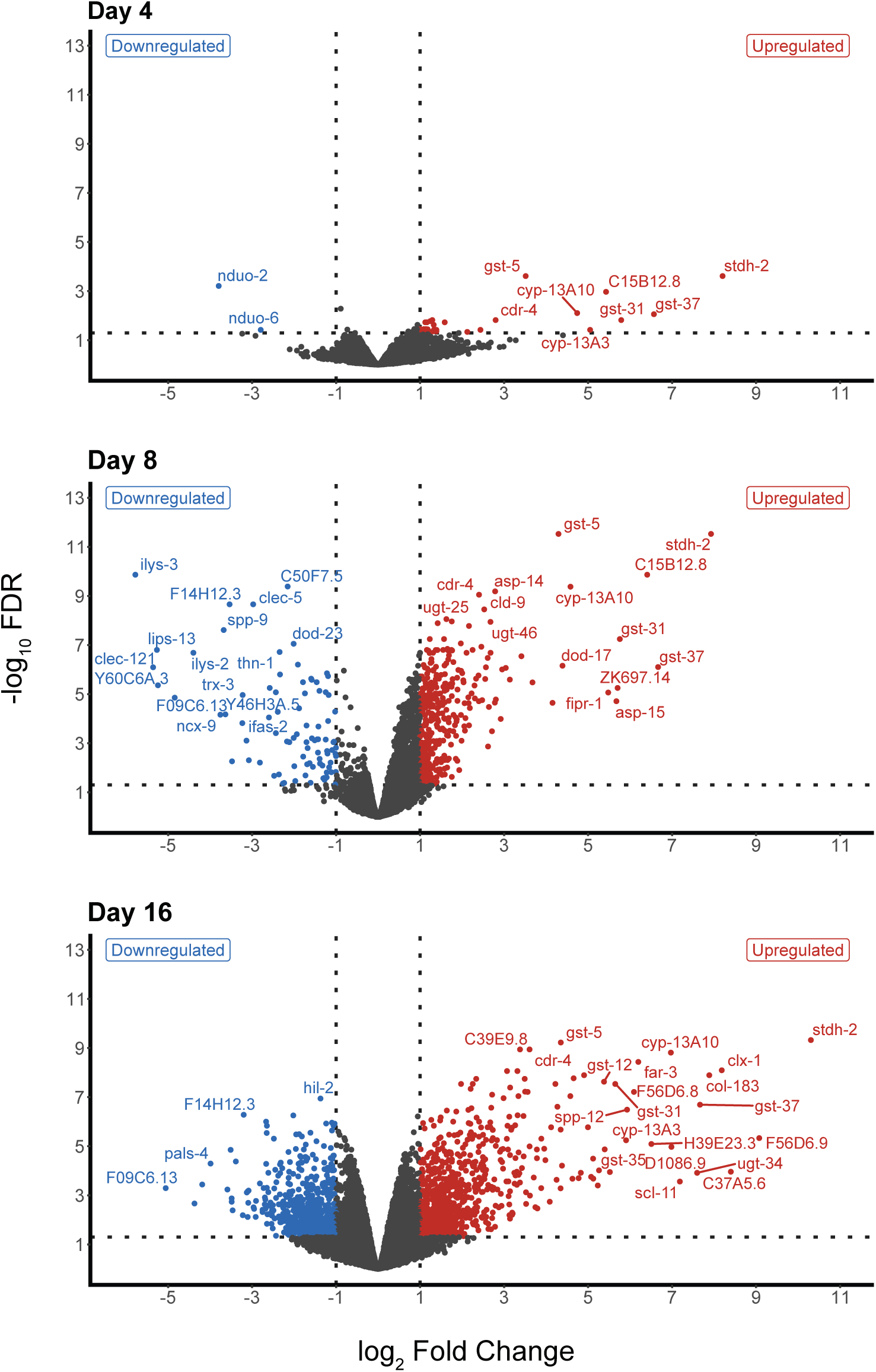
Volcano plot of differentially expressed genes (DEGs) at days 4, 8, and 16 of sulforaphane treatment. Significantly DE genes with absolute log_2_ fold change of at least 1 and FDR of 0.05 or lower are colored blue (downregulated) and red (upregulated).

**Extended Data Fig. 3:**
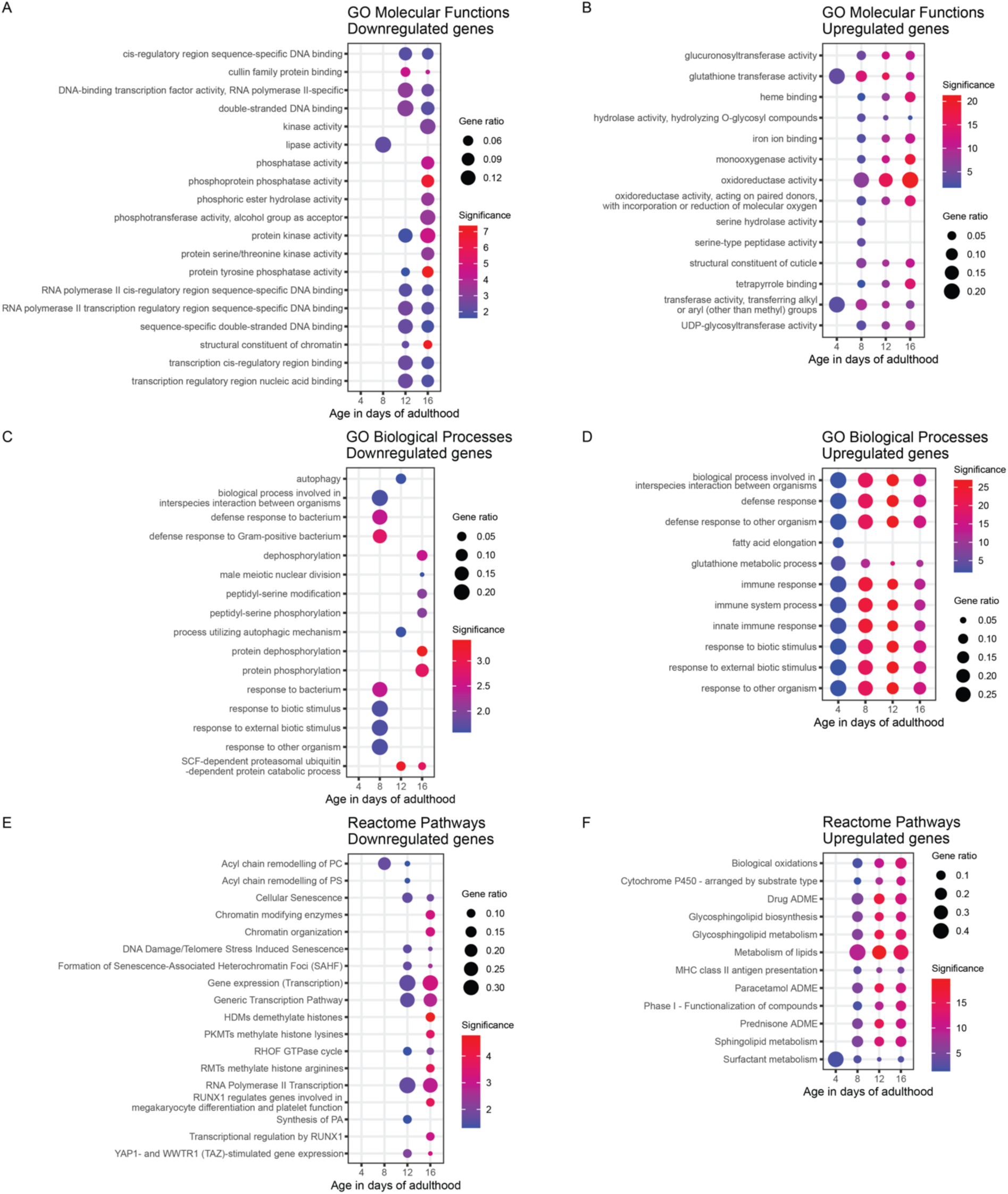
Gene Ontology molecular functions (A, downregulated; B, upregulated), Gene Ontology biological processes (C, downregulated; D, upregulated) and Reactome pathway (E, downregulated; D, upregulated) enrichments after treatment with 100 uM sulforaphane at ages 4, 8, 12, and 16 of adulthood.

**Extended Data Fig. 4:**
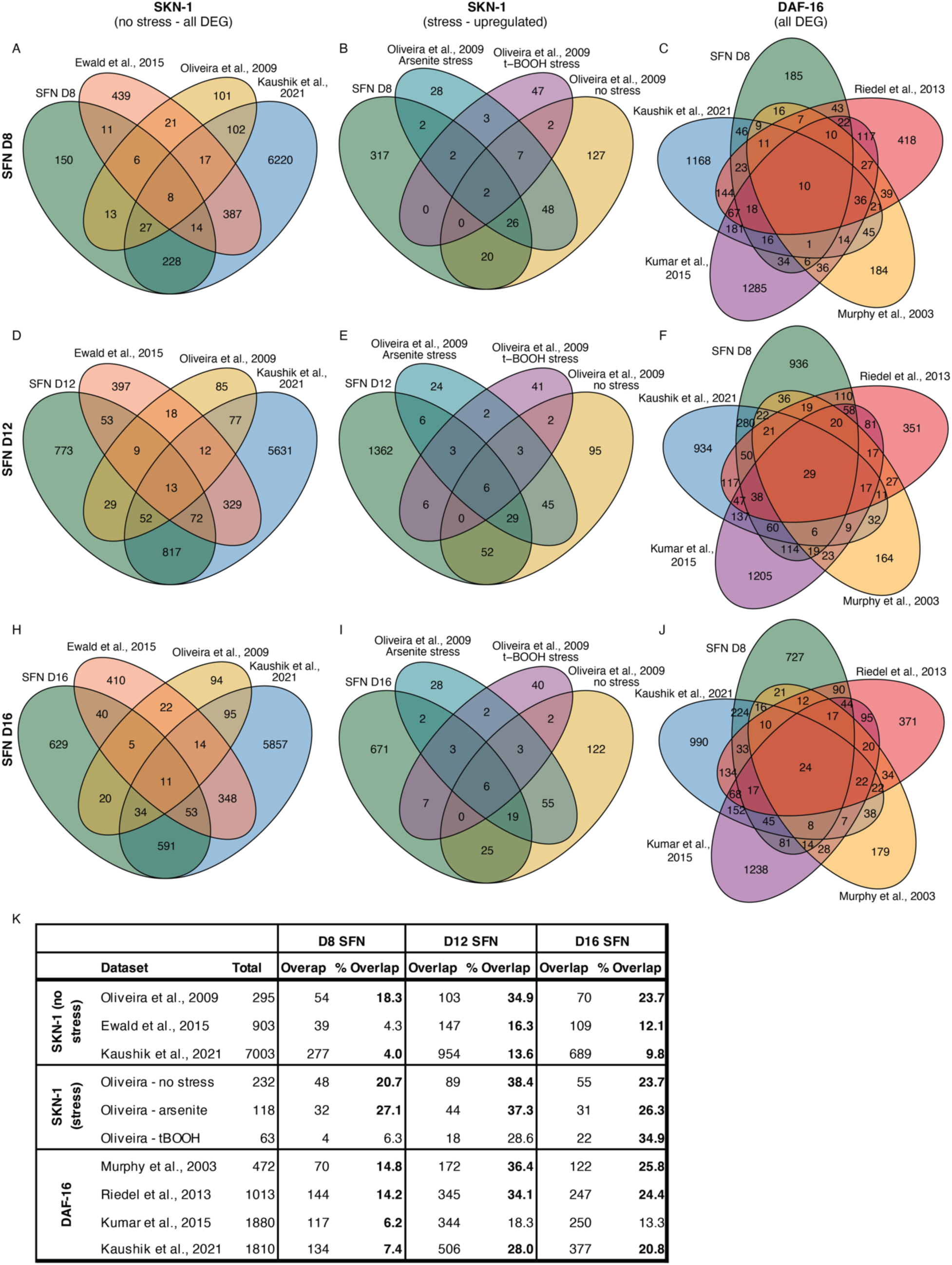
Venn diagrams showing the overlap of differentially expressed genes with SFN treatment to various mutant expression datasets from the literature. Comparison of differentially expressed SFN genes at day 8 (A-C), day 12 (D-F), and day 16 (H-J) to published SKN-1 and DAF-16 RNA-seq and microarray datasets. Venn diagrams show the overlap of our SFN DEG to published SKN-1 datasets (all DEG) with stress (A, D and H), and upregulated genes only under arsenite or t-BOOH stress (B, E, and I). Venn diagrams comparing SFN DEG to published DAF-16 target datasets (C, F, and J). (K) Summary table of the overlap and percent overlap of all datasets references with our SFN data at various ages. Overlap percentages where *p<*0.05 with a Fisher’s exact test are denoted in bold.

**Supplementary Table 1:** The transcriptomic response to sulforaphane is delayed and increases with age. The total number of differentially expressed genes (DEGs) at each age collected for RNA-seq. Genes with an absolute log_2_ fold change of at least 1 and FDR of 0.05 or lower are significantly differentially expressed. All samples started treatment with 100 uM sulforaphane on day 1 of adulthood.

| Age (day of adulthood) | Total DEGs | Upregulated | Downregulated | No significant change | Average reads from 4 samples (millions) control/SFN |
| --- | --- | --- | --- | --- | --- |
| 2 | 12 | 12 | 0 | 9390 | 70.6/45.1 |
| 4 | 31 | 29 | 2 | 9196 | 31.6/13.7 |
| 8 | 457 | 369 | 88 | 10298 | 51.4/48.2 |
| 12 | 1818 | 1464 | 354 | 10904 | 55.6/43.8 |
| 16 | 1383 | 733 | 650 | 10453 | 45.3/46.3 |

**Supplementary Table 2:** Detoxification genes are significantly upregulated compared to the entire transcriptome. The number of genes identified in total, the number upregulated, and the percent upregulated for the entire transcriptome (overall), the CYP, GST, and UGT gene families at ages 2, 4, 12, and 16 of adulthood. Upregulation refers to genes with a log fold change of >1 and false discovery rate <.05 with 100 uM SFN treatment as compared to the DMSO control. *p*-values indicate significance with a Fischer’s exact test.

| Sample age | Gene family | Total genes | Upregulated genes | % Upregulated | p value |
| --- | --- | --- | --- | --- | --- |
| 2 | Overall | 9402 | 12 | 0.1 |  |
| 2 | CYP | 37 | 1 | 2.7 | 0.04606 |
| 2 | GST | 25 | 2 | 8.0 | 0.0004385 |
| 2 | UGT | 46 | 0 | 0.0 | 1 |
| 4 | Overall | 9227 | 29 | 0.3 |  |
| 4 | CYP | 36 | 2 | 5.6 | 0.005582 |
| 4 | GST | 25 | 4 | 16.0 | 9.41E-07 |
| 4 | UGT | 42 | 1 | 2.4 | 0.1236 |
| 8 | Overall | 10755 | 369 | 3.4 |  |
| 8 | CYP | 44 | 8 | 18.2 | 0.0001038 |
| 8 | GST | 30 | 16 | 53.3 | 2.404E-16 |
| 8 | UGT | 46 | 11 | 23.9 | 2.904E-07 |
| 12 | Overall | 12722 | 1464 | 11.5 |  |
| 12 | CYP | 46 | 26 | 56.5 | 1.62E-13 |
| 12 | GST | 36 | 26 | 72.2 | < 2.2e-16 |
| 12 | UGT | 53 | 32 | 60.4 | 2.2E-16 |
| 16 | Overall | 11836 | 733 | 6.2 |  |
| 16 | CYP | 45 | 25 | 55.6 | < 2.2e-16 |
| 16 | GST | 28 | 17 | 60.7 | 2.63E-14 |
| 16 | UGT | 47 | 22 | 46.8 | 6.09E-15 |

**Supplementary Table 3:**
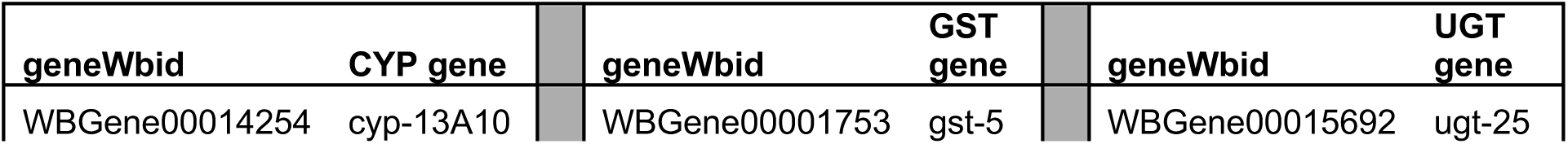
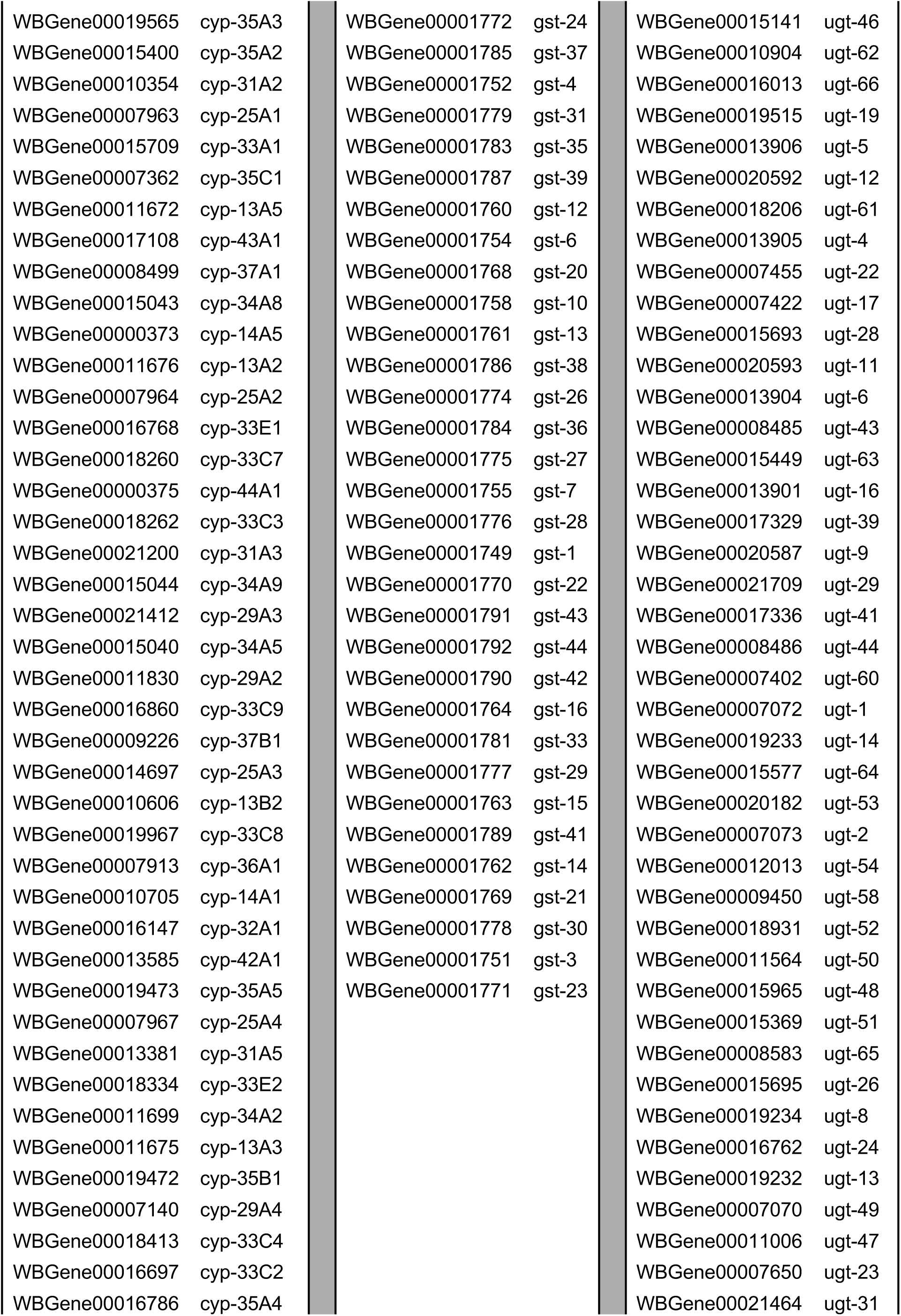
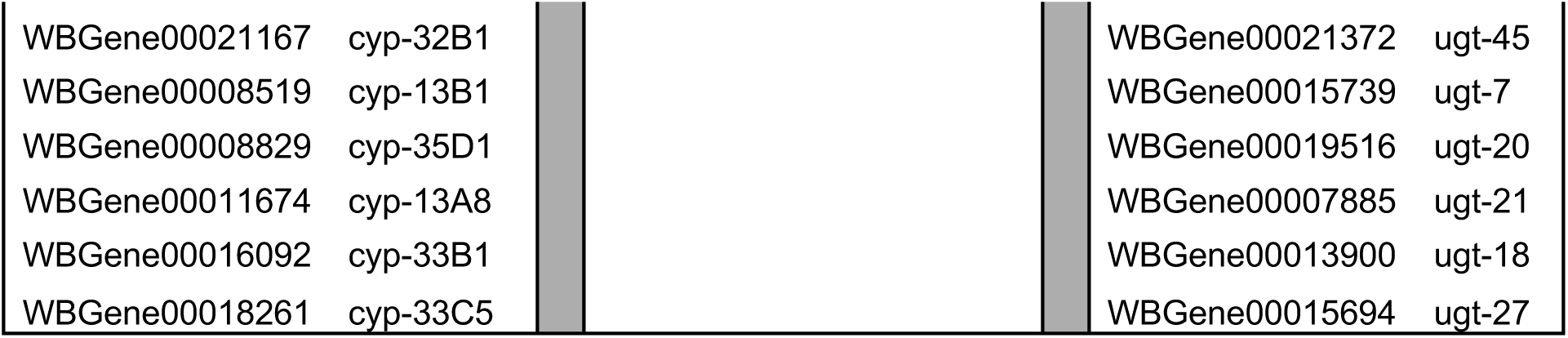
List of all detoxification genes in Figure 4.

## Notes

### Competing Interest Statement

The authors have declared no competing interest.

