## Supplemental figure 1 for "The broccoli derivative sulforaphane extends lifespan by slowing the transcriptional aging clock"

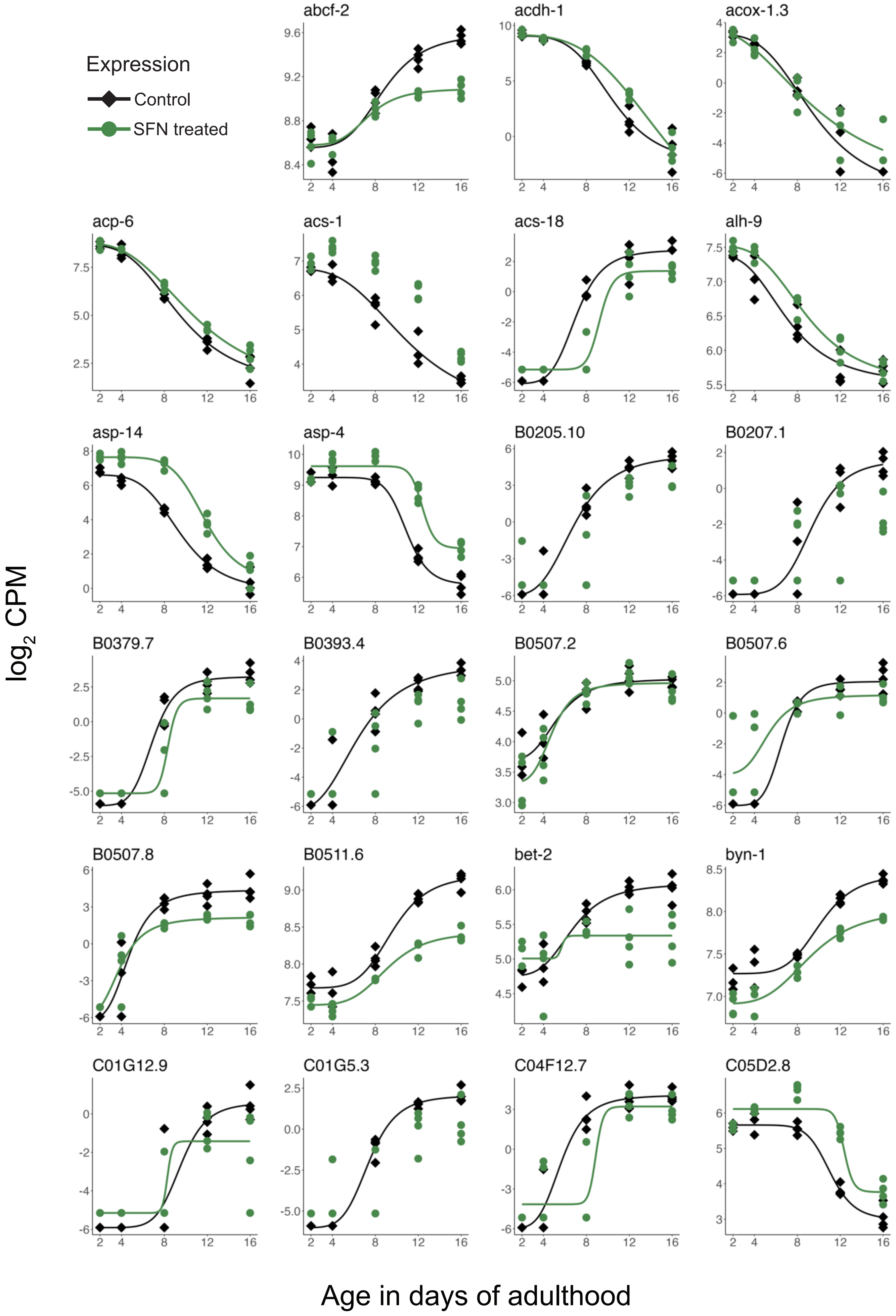
Expression
Control
SFN treated
abcf-2
acdh-1
acox-1.3
acp-6
acs-1
acs-18
alh-9
asp-14
asp-4
B0205.10
B0207.1
B0379.7
B0393.4
B0507.2
B0507.6
B0507.8
B0511.6
bet-2
byn-1
C01G12.9
C01G5.3
C04F12.7
C05D2.8
log2 CPM
Age in days of adulthood

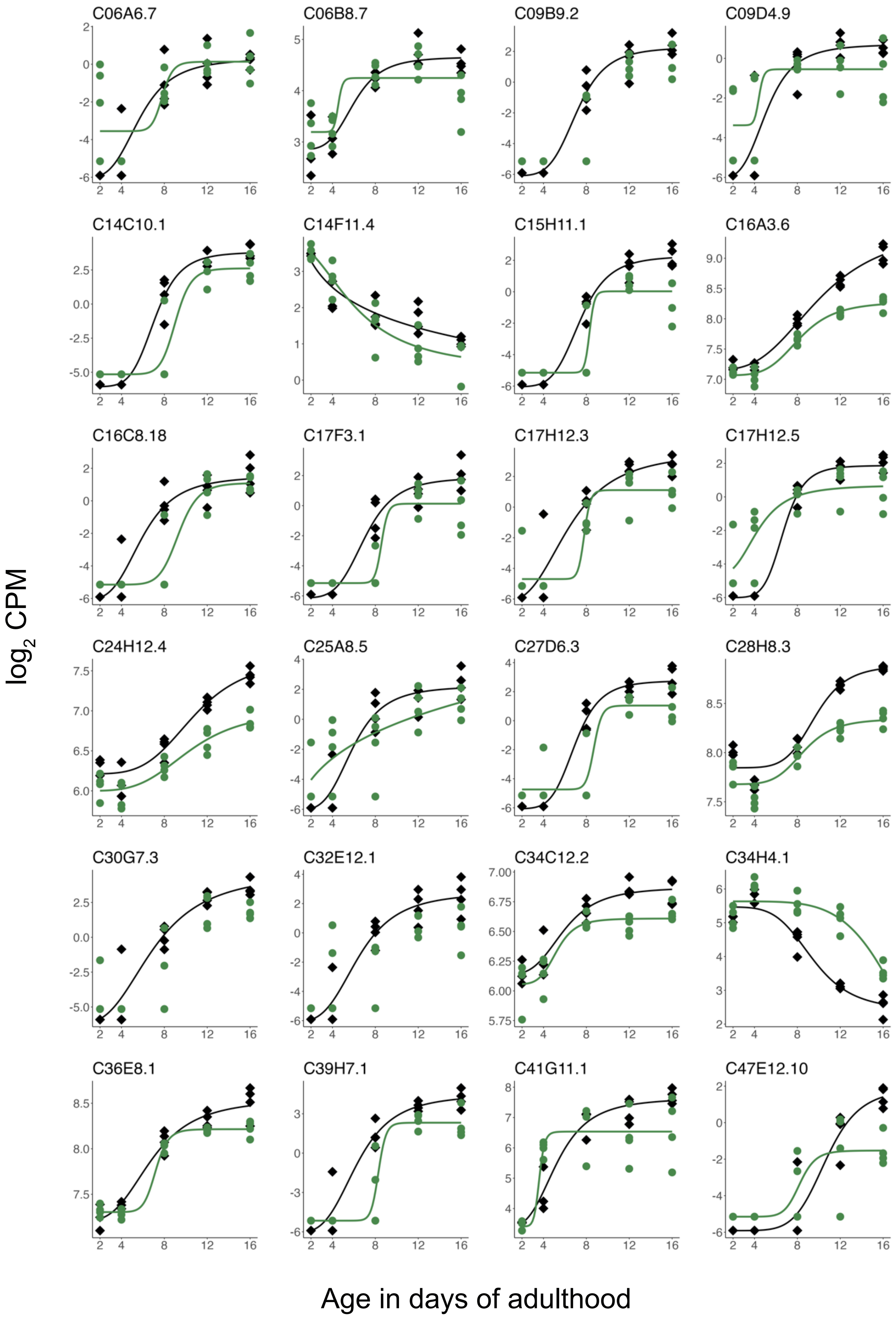
C06A6.7
C06B8.7
C09B9.2
C09D4.9
C14C10.1
C14F11.4
C15H11.1
C16A3.6
C16C8.18
C17F3.1
C17H12.3
C17H12.5
C24H12.4
C25A8.5
C27D6.3
C28H8.3
C30G7.3
C32E12.1
C34C12.2
C34H4.1
C36E8.1
C39H7.1
C41G11.1
C47E12.10
$\log_2$ CPM
Age in days of adulthood

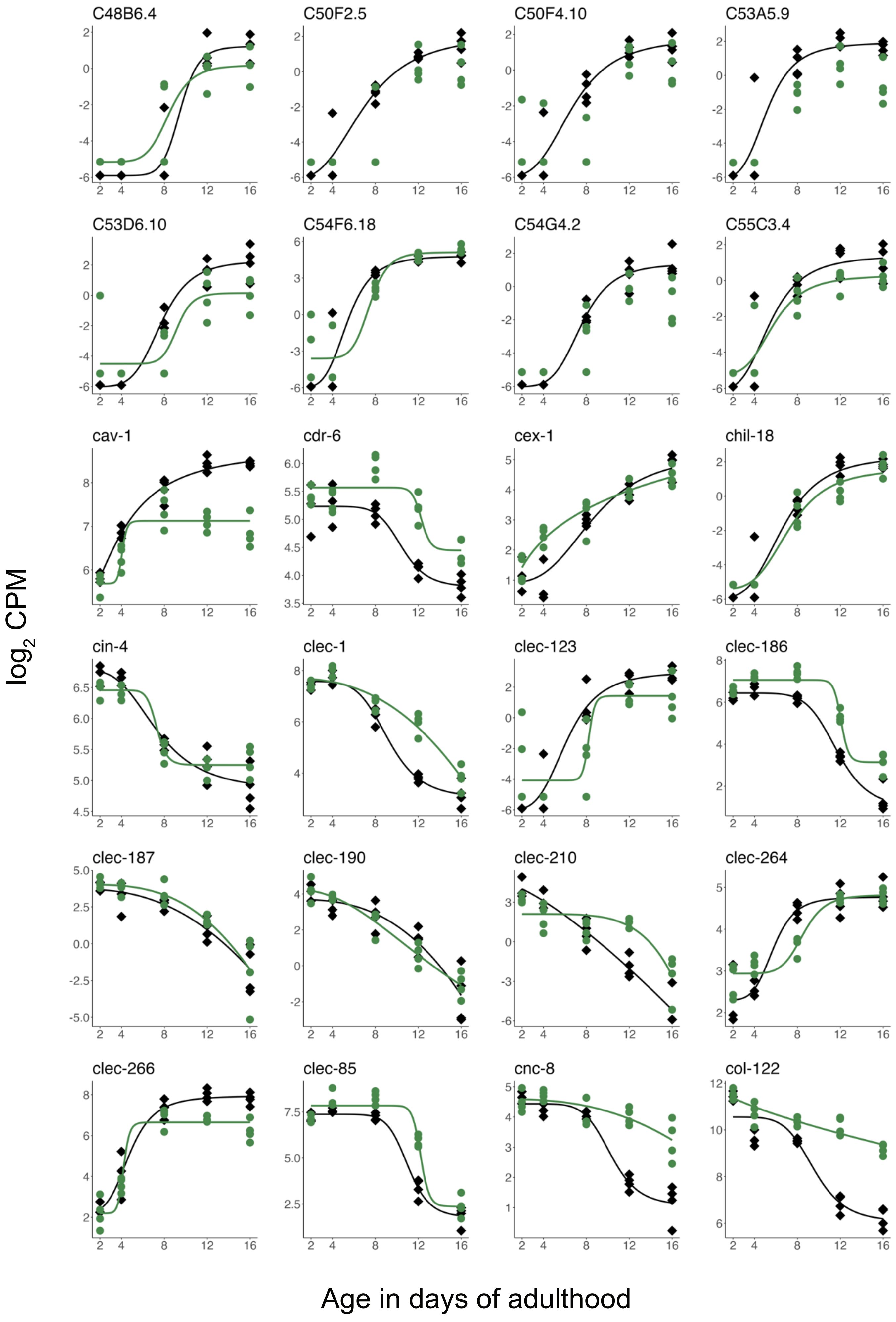
C48B6.4
C50F2.5
C50F4.10
C53A5.9
C53D6.10
C54F6.18
C54G4.2
C55C3.4
cav-1
cdr-6
cex-1
chil-18
cin-4
clec-1
clec-123
clec-186
clec-187
clec-190
clec-210
clec-264
clec-266
clec-85
cnc-8
col-122
$\log_2$ CPM
Age in days of adulthood

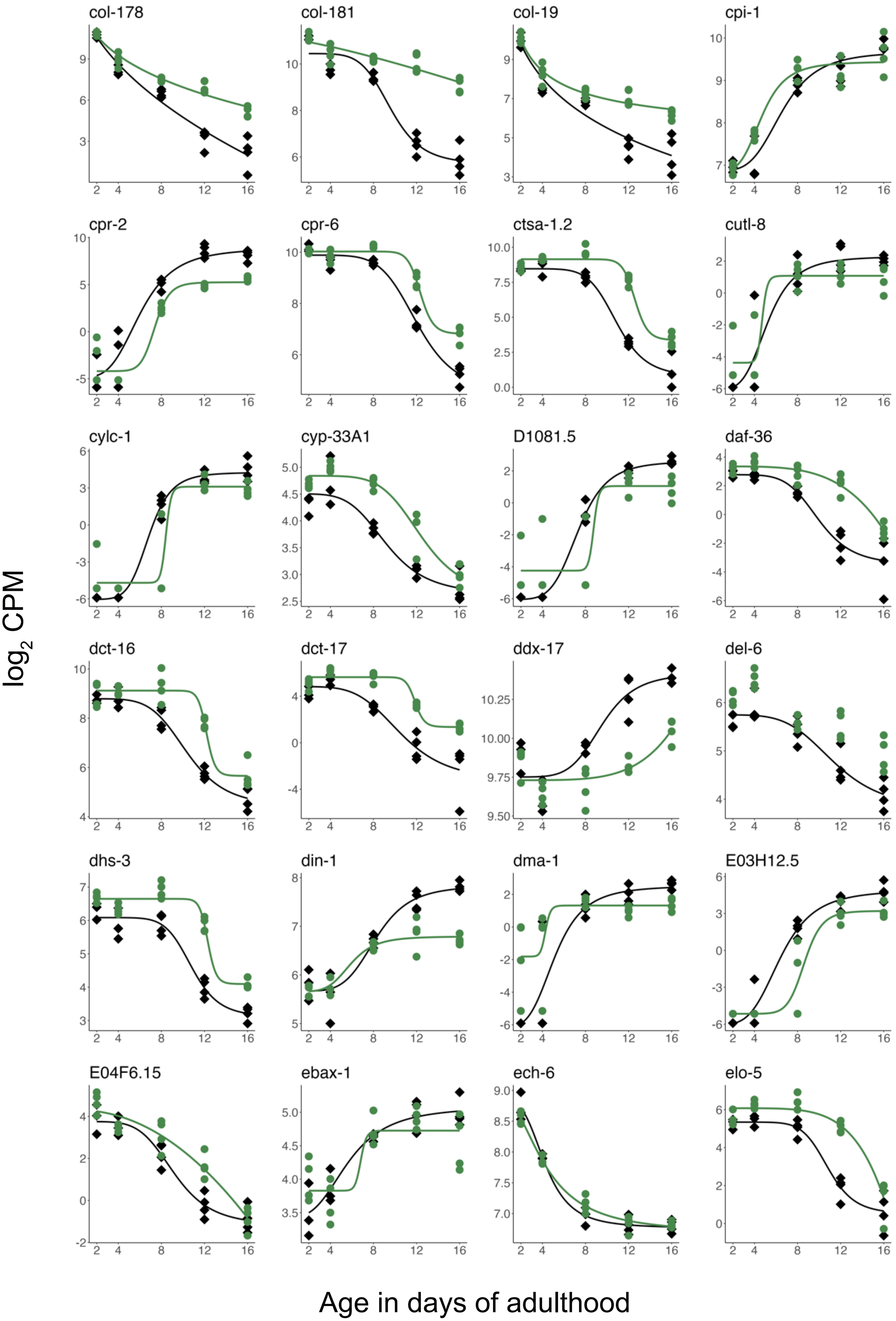
col-178
col-181
col-19
cpi-1
cpr-2
cpr-6
ctsa-1.2
cutl-8
cylc-1
cyp-33A1
D1081.5
daf-36
dct-16
dct-17
ddx-17
del-6
dhs-3
din-1
dma-1
E03H12.5
E04F6.15
ebax-1
ech-6
elo-5
$\log_2$ CPM
Age in days of adulthood

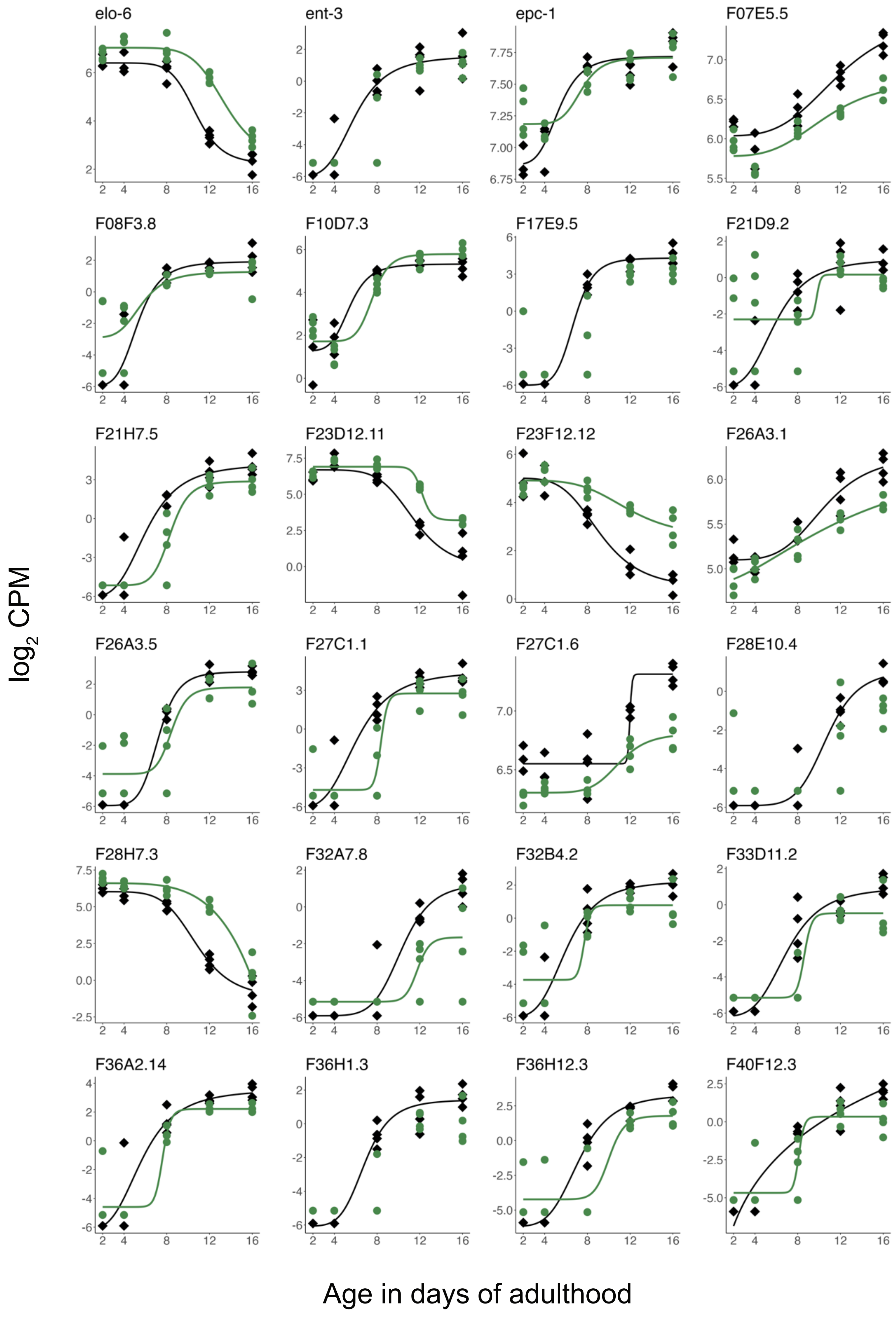

elo-6
ent-3
epc-1
F07E5.5
F08F3.8
F10D7.3
F17E9.5
F21D9.2
F21H7.5
F23D12.11
F23F12.12
F26A3.1
F26A3.5
F27C1.1
F27C1.6
F28E10.4
F28H7.3
F32A7.8
F32B4.2
F33D11.2
F36A2.14
F36H1.3
F36H12.3
F40F12.3
log2 CPM
Age in days of adulthood

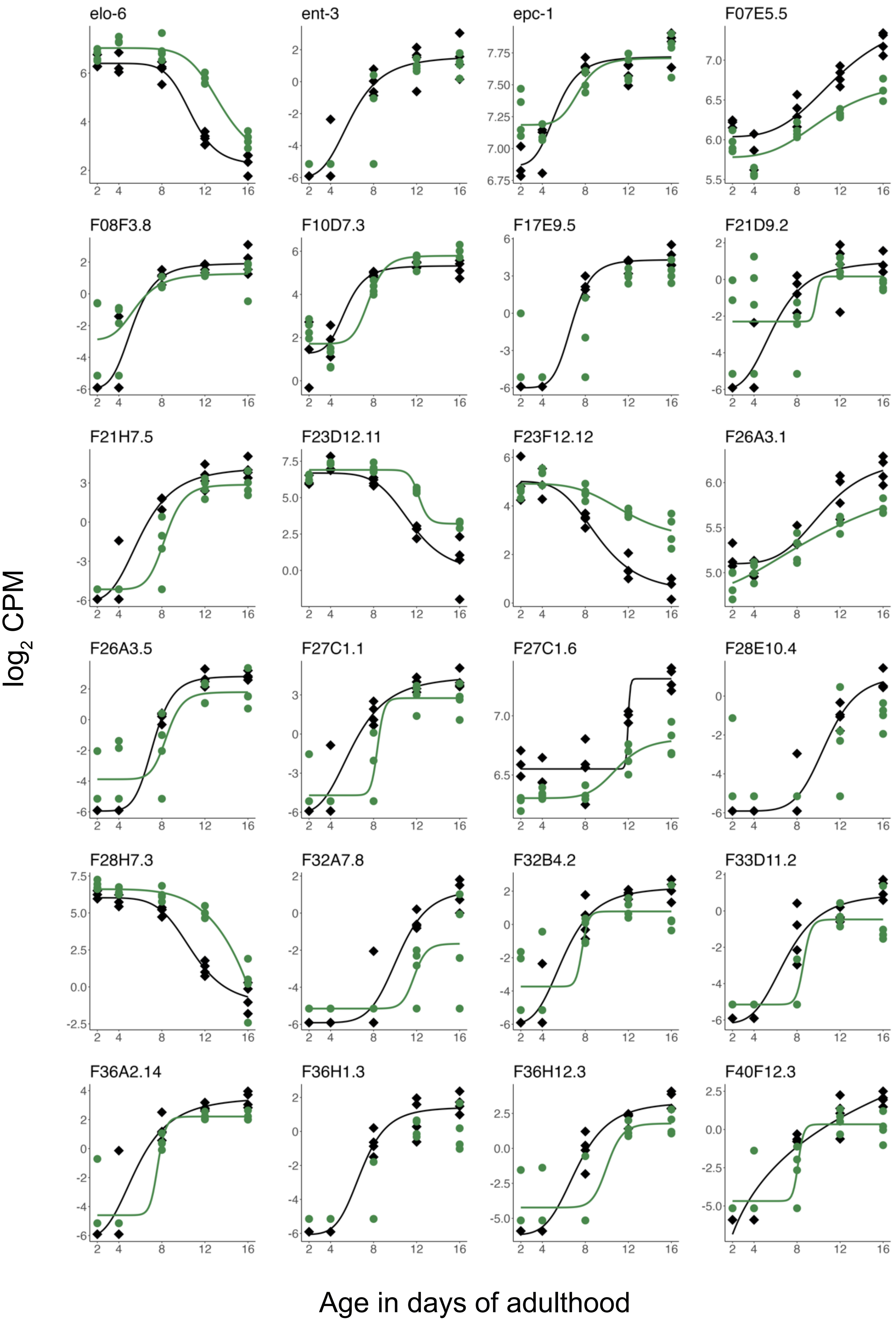

elo-6
ent-3
epc-1
F07E5.5
F08F3.8
F10D7.3
F17E9.5
F21D9.2
F21H7.5
F23D12.11
F23F12.12
F26A3.1
F26A3.5
F27C1.1
F27C1.6
F28E10.4
F28H7.3
F32A7.8
F32B4.2
F33D11.2
F36A2.14
F36H1.3
F36H12.3
F40F12.3
$\log_2$ CPM
Age in days of adulthood

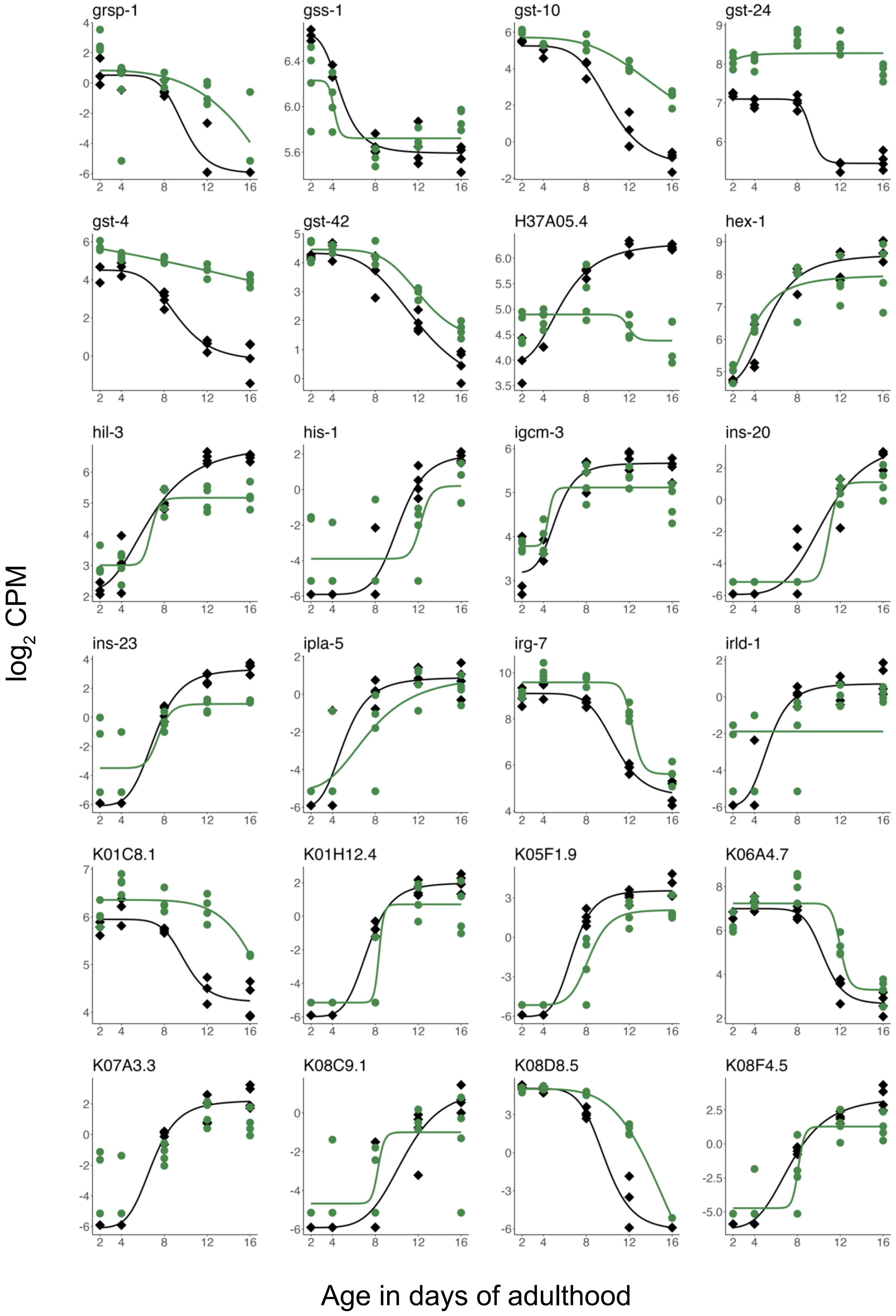
grsp-1
gss-1
gst-10
gst-24
gst-4
gst-42
H37A05.4
hex-1
hil-3
his-1
igcm-3
ins-20
ins-23
ipla-5
irg-7
irld-1
K01C8.1
K01H12.4
K05F1.9
K06A4.7
K07A3.3
K08C9.1
K08D8.5
K08F4.5
$\log_2$ CPM
Age in days of adulthood

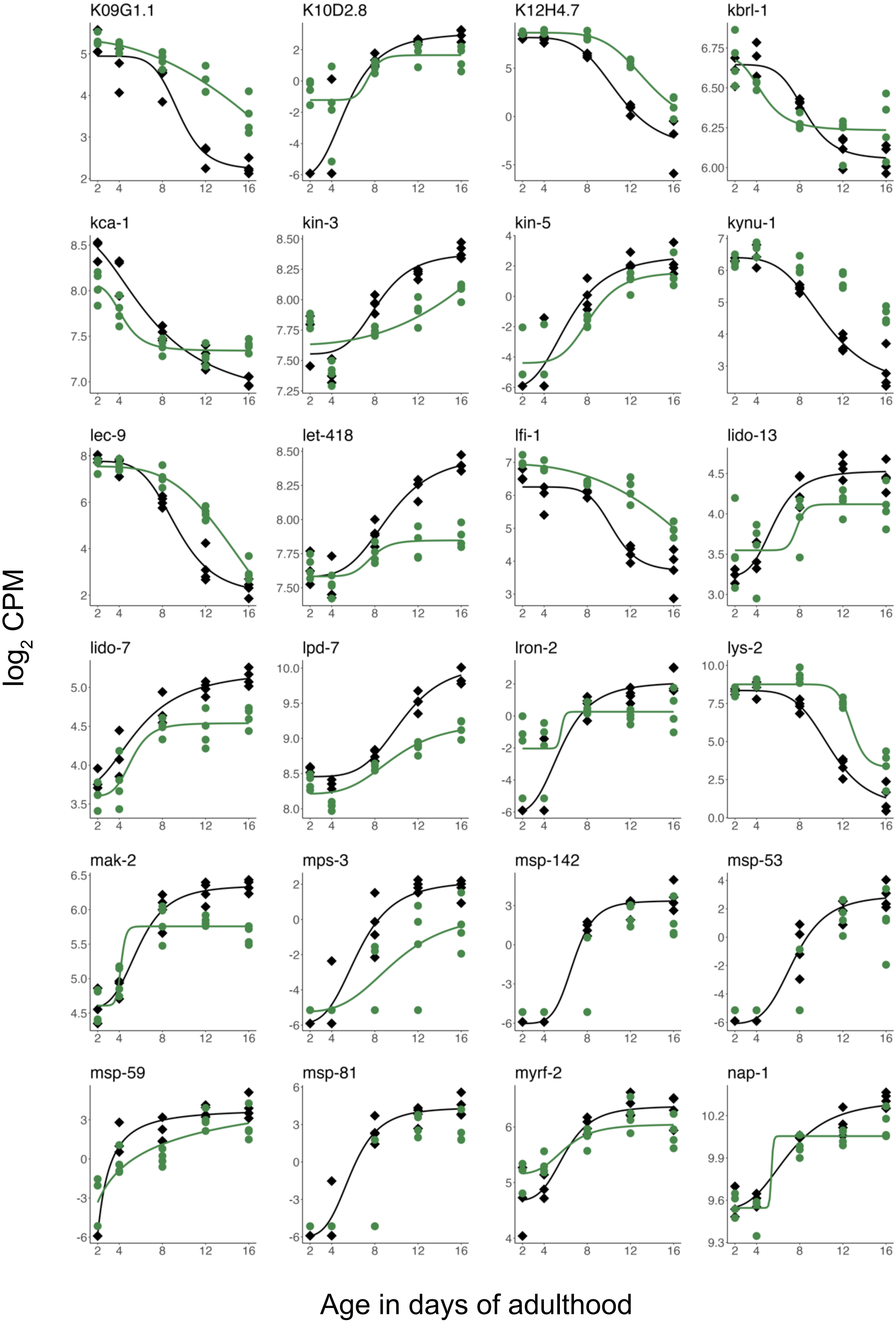
K09G1.1
K10D2.8
K12H4.7
kbrl-1
kca-1
kin-3
kin-5
kynu-1
lec-9
let-418
lfi-1
lido-13
lido-7
lpd-7
lron-2
lys-2
mak-2
mps-3
msp-142
msp-53
msp-59
msp-81
myrf-2
nap-1
$\log_2$ CPM
Age in days of adulthood

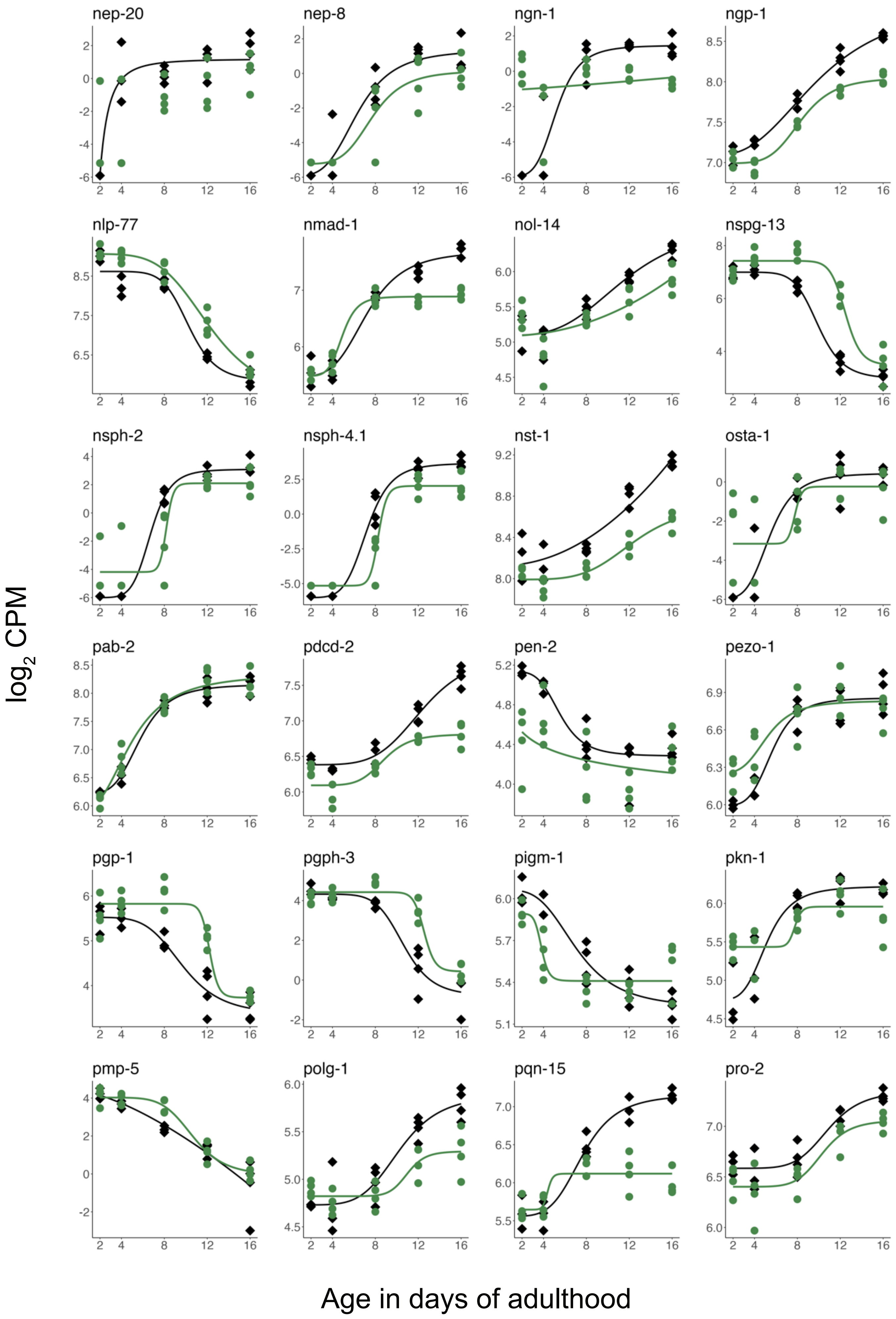
nep-20
nep-8
ngn-1
ngp-1
nlp-77
nmad-1
nol-14
nspg-13
nsph-2
nsph-4.1
nst-1
osta-1
pab-2
pdcd-2
pen-2
pezo-1
pgp-1
pgph-3
pigm-1
pkn-1
pmp-5
polg-1
pqn-15
pro-2
$\log_2$ CPM
Age in days of adulthood

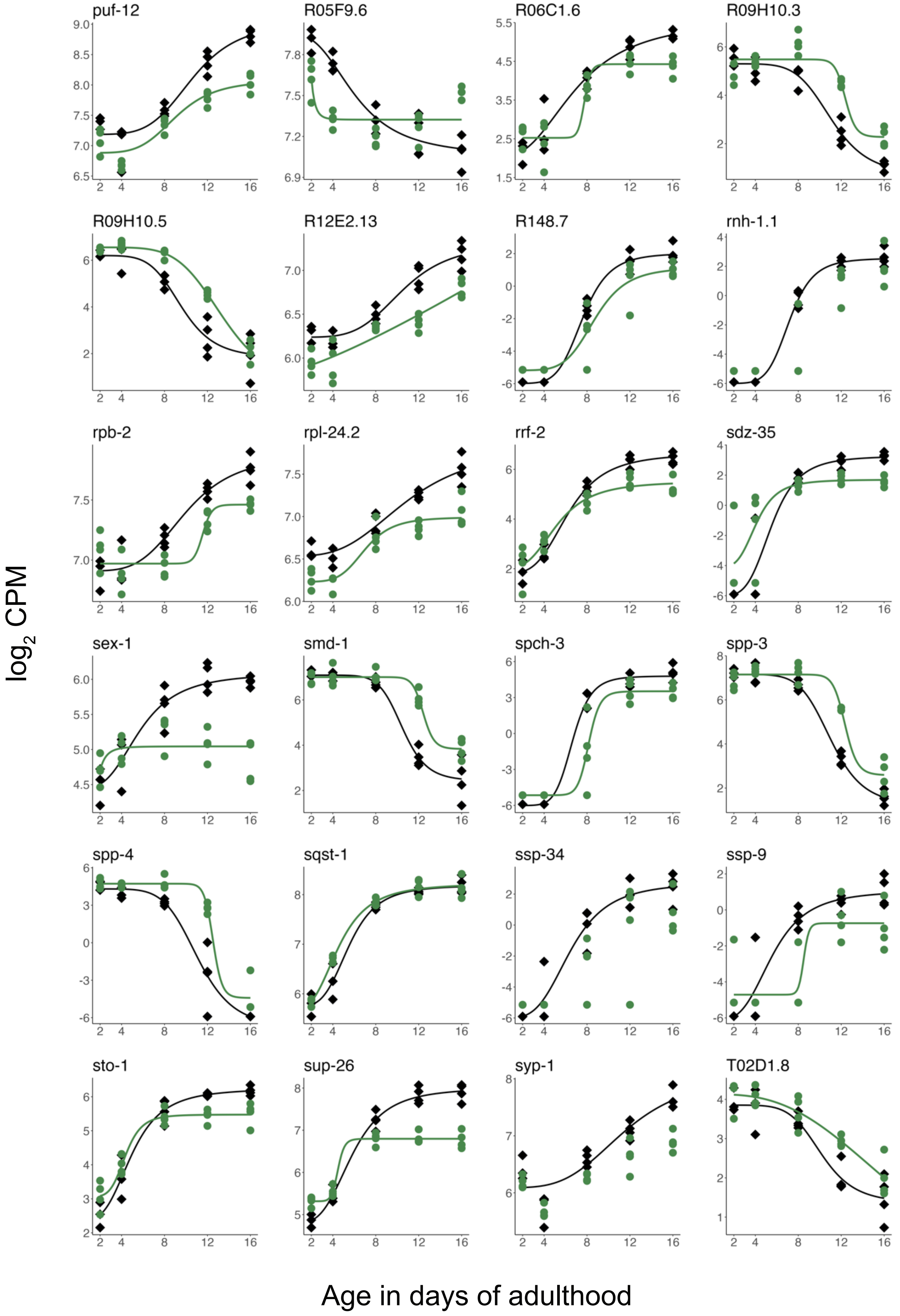

puf-12
R05F9.6
R06C1.6
R09H10.3
R09H10.5
R12E2.13
R148.7
rnh-1.1
rpb-2
rpl-24.2
rrf-2
sdz-35
sex-1
smd-1
spch-3
spp-3
spp-4
sqst-1
ssp-34
ssp-9
sto-1
sup-26
syp-1
T02D1.8
log2 CPM
Age in days of adulthood

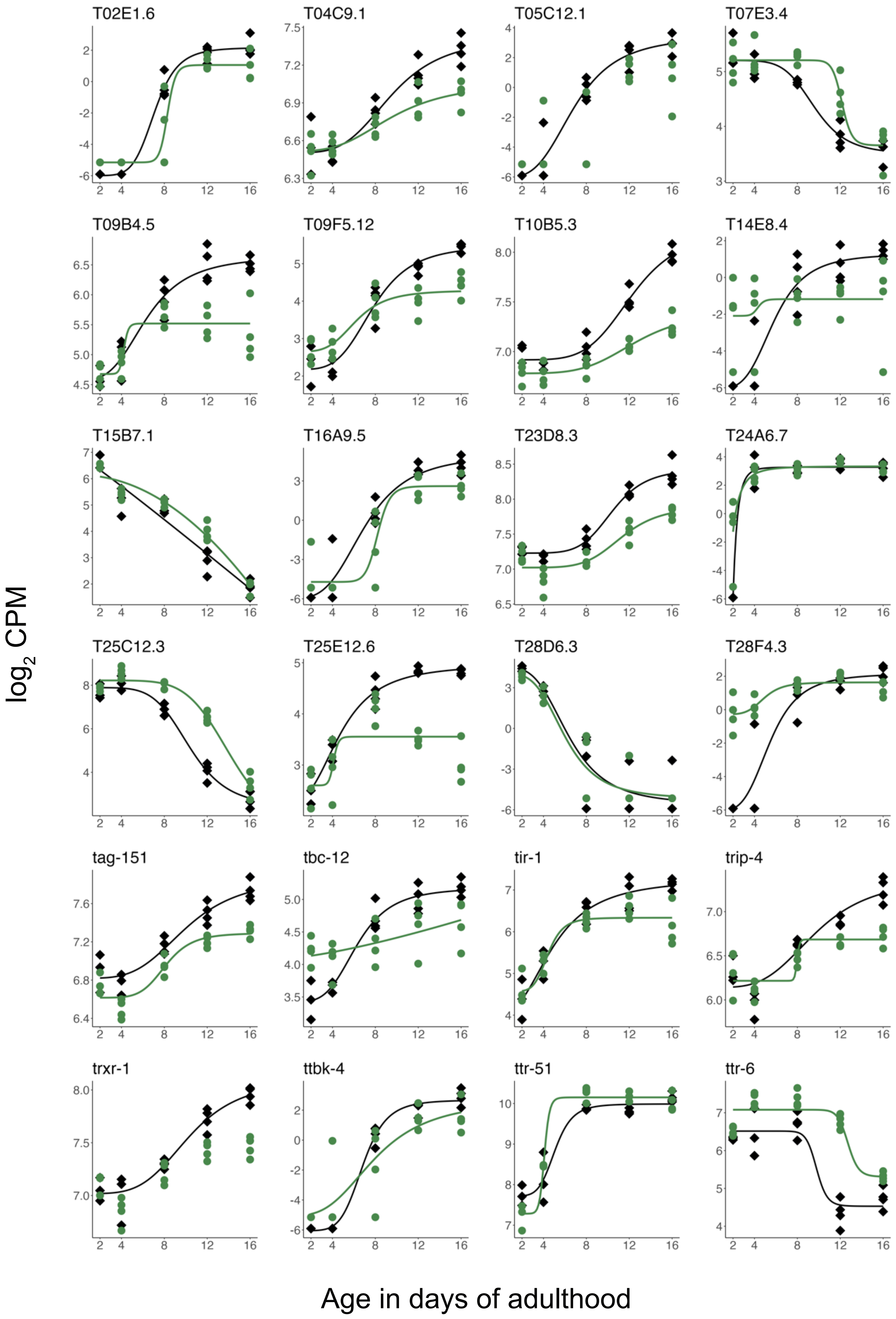

T02E1.6
T04C9.1
T05C12.1
T07E3.4
T09B4.5
T09F5.12
T10B5.3
T14E8.4
T15B7.1
T16A9.5
T23D8.3
T24A6.7
T25C12.3
T25E12.6
T28D6.3
T28F4.3
tag-151
tbc-12
tir-1
trip-4
trxr-1
ttbk-4
ttr-51
ttr-6
$\log_2$ CPM
Age in days of adulthood

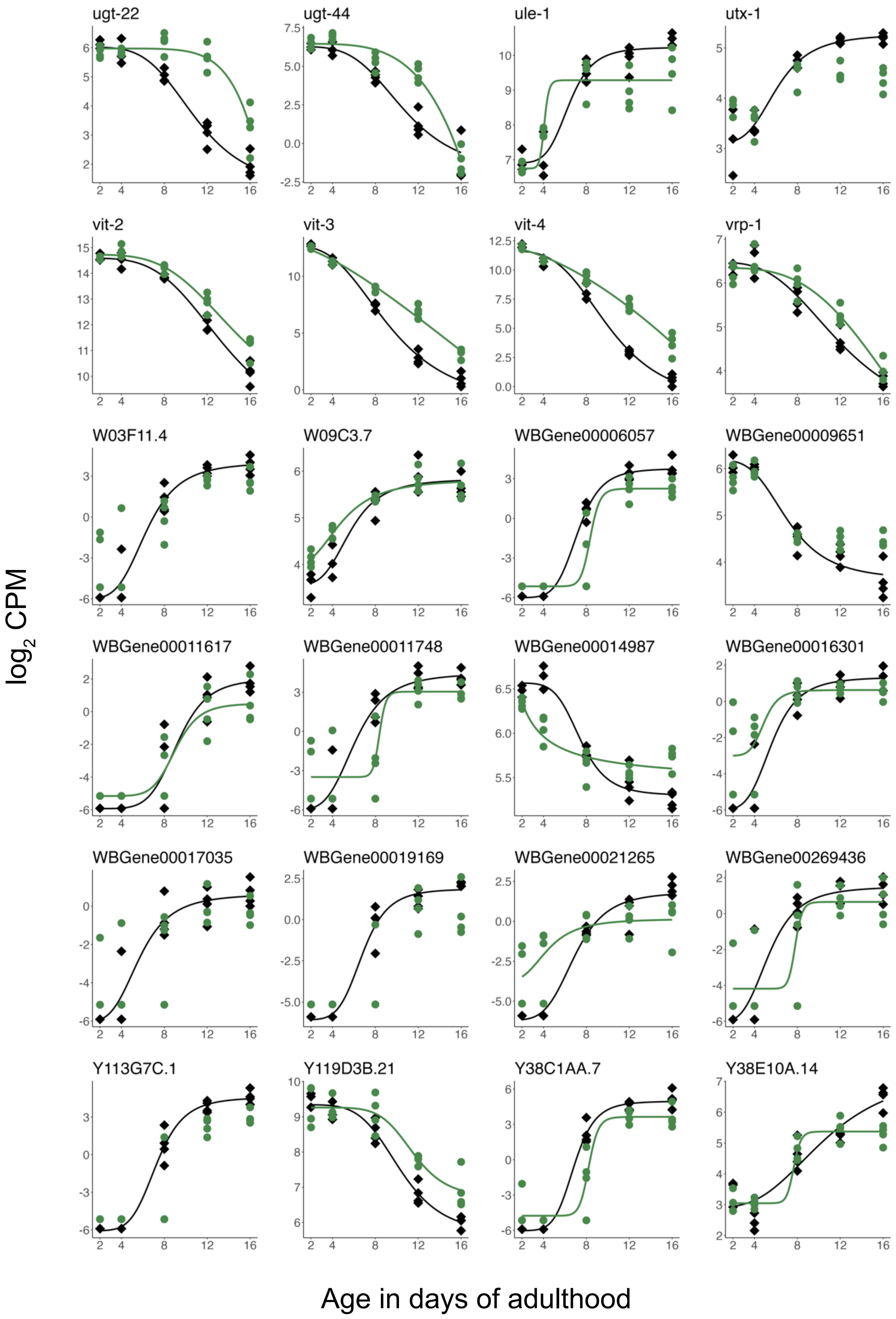
ugt-22
ugt-44
ule-1
utx-1
vit-2
vit-3
vit-4
vrp-1
W03F11.4
W09C3.7
WBGene00006057
WBGene00009651
WBGene00011617
WBGene00011748
WBGene00014987
WBGene00016301
WBGene00017035
WBGene00019169
WBGene00021265
WBGene00269436
Y113G7C.1
Y119D3B.21
Y38C1AA.7
Y38E10A.14
$\log_2$ CPM
Age in days of adulthood

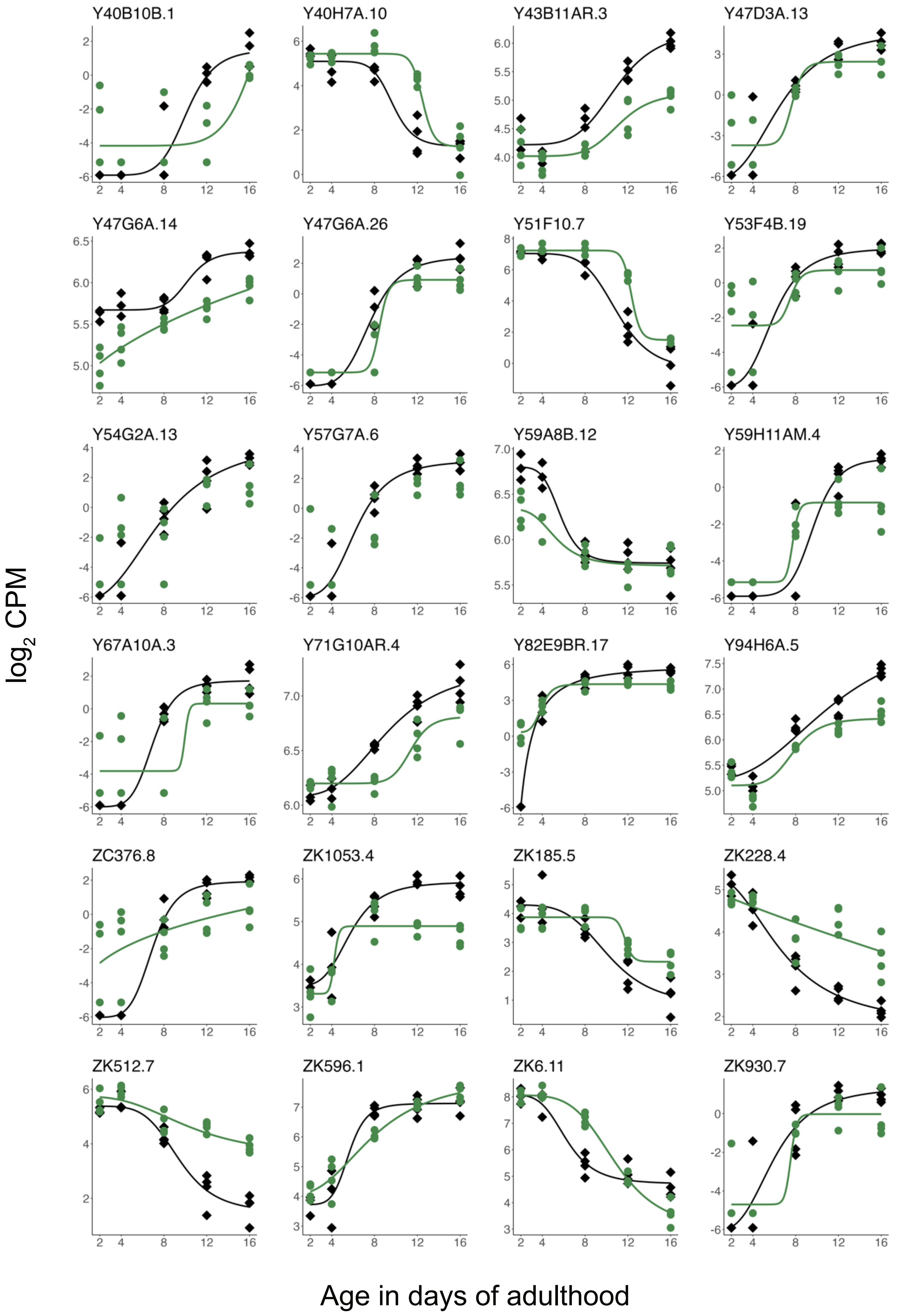
Y40B10B.1
Y40H7A.10
Y43B11AR.3
Y47D3A.13
Y47G6A.14
Y47G6A.26
Y51F10.7
Y53F4B.19
Y54G2A.13
Y57G7A.6
Y59A8B.12
Y59H11AM.4
Y67A10A.3
Y71G10AR.4
Y82E9BR.17
Y94H6A.5
ZC376.8
ZK1053.4
ZK185.5
ZK228.4
ZK512.7
ZK596.1
ZK6.11
ZK930.7
log2 CPM
Age in days of adulthood
